# *Chlamydomonas* ARMC2/PF27 is an obligate cargo adapter for IFT of radial spokes

**DOI:** 10.1101/2021.10.31.466660

**Authors:** Karl Lechtreck, Yi Liu, Jin Dai, Rama A. Alkhofash, Jack Butler, Lea Alford, Pinfen Yang

## Abstract

Intraflagellar transport (IFT) carries proteins into flagella but how IFT trains interact with the large number of diverse proteins required to assemble flagella remains largely unknown. Here, we show that IFT of radial spokes in *Chlamydomonas* requires ARMC2/PF27, a conserved armadillo repeat protein associated with male infertility and reduced lung function. *Chlamydomonas* ARMC2 was highly enriched in growing flagella and tagged ARMC2 and the spoke protein RSP3 comigrated on anterograde trains. In contrast, a cargo and an adapter of inner and outer dynein arms moved independently of ARMC2, indicating that unrelated cargoes distribute stochastically onto the IFT trains. After concomitant unloading at the flagellar tip, RSP3 attached to the axoneme whereas ARMC2 diffused back to the cell body. In *armc2/pf27* mutants, IFT of radial spokes was abolished and the presence of radial spokes was limited to the proximal region of flagella. We conclude that ARMC2 is a cargo adapter required for IFT of radial spokes to ensure their assembly along flagella. ARMC2 belongs to a growing class of cargo-specific adapters that enable flagellar transport of preassembled axonemal substructures by IFT.

## Introduction

Cilia and eukaryotic flagella consist of hundreds of distinct proteins, which are synthesized in the cell body and moved posttranslationally into the organelle (Rosenbaum and Child 1967; Pazour *et al*. 2005). Protein transport into flagella involves intraflagellar transport (IFT), a motor-based bidirectional motility of protein carriers (i.e., “IFT trains”)(Kozminski *et al*. 1993). Numerous proteins of the flagellar axoneme, matrix, and membrane have been shown to use the IFT pathway for flagellar entry and exit (Lechtreck 2015). This raises the question how IFT trains, composed of just 22 IFT proteins and the associated kinesin-2 and IFT dynein motors, interact with the large number of diverse flagellar proteins. Tubulin, the most abundant flagellar protein, interacts directly with the N-terminal domains of the IFT-B core proteins IFT74 and IFT81 (Bhogaraju *et al*. 2013; Craft *et al*. 2015; Kubo *et al*. 2016; Craft Van De Weghe *et al*. 2020). However, many other proteins do not bind directly to the IFT trains but interaction is mediated by IFT cargo adapters. The octameric BBSome, for example, acts as a linker for a diverse group of transmembrane and membrane-associated proteins, attaching them indirectly to IFT trains (Nachury *et al*. 2007; Liu and Lechtreck 2018; Wingfield *et al*. 2018). With respect to axonemal proteins, IFT of outer dynein arms (ODAs) and the inner dynein arm I1/f requires the adapter proteins ODA16 and IDA3, respectively (Ahmed *et al*. 2008; Dai *et al*. 2018; Hunter *et al*. 2018). The corresponding *oda16* and *ida3* mutants assemble full-length flagella that specifically lack ODAs or IDAs I1/f, respectively, but of otherwise normal ultrastructure (Ahmed and Mitchell 2005; Hunter *et al*. 2018). ODAs and IDAs are large multiprotein complexes, which are assembled in the cell body before the entire substructures are moved into the flagella by IFT (Fowkes and Mitchell 1998; King 2012; Viswanadha *et al*. 2014). Similarly, more than 20 radial spoke (RS) proteins preassemble into a 12S RS precursor in the cell body (Qin *et al*. 2004; Yang *et al*. 2006). Then, the L-shaped precursors are moved by IFT to the flagellar tip, converted into the mature 20S spoke complexes, and assembled as T-shaped spokes onto the axonemal doublets (Qin *et al*. 2004; Diener *et al*. 2011; Lechtreck *et al*. 2018; GROSSMAN-HAHAM *et al*. 2021; Gui *et al*. 2021). Mutations in the genes encoding the various spoke subunits lead to partial or complete loss of the RSs and flagellar paralysis (Luck *et al*. 1977; Piperno *et al*. 1977; Witman *et al*. 1978; Piperno *et al*. 1981). The *pf27* mutant, however, stands out as it assembles radial spokes of normal ultrastructure and subunit composition but the presence of spokes is limited to the very proximal region of the mutant flagella (Huang *et al*. 1981; Alford *et al*. 2013). *In vitro* decoration experiments using isolated axonemes and RSs revealed that the *pf27* axonemes bind control and *pf27* spokes, indicating that axonemal docking of RSs is unaffected in *pf27* (Alford *et al*. 2013). To explain the absence of spokes from large sections of the *pf27* flagella, Alford et al. (2013) postulated that *PF27* could encode a factor required for the transport of radial spokes into the distal flagellum via IFT. Then, RS assembly in the proximal region of *pf27* flagella could result from residual entry of RSs by diffusion followed by binding to the nearest available docking sites. Such a scenario could also explain why the phosphorylation state of several RS proteins is altered in *pf27* as it has been proposed that phosphorylation of RSs occurs near the flagellar tip, which the RSs would fail to reach in *pf27* (Huang *et al*. 1981; Yang and Yang 2006; Gupta *et al*. 2012). The *pf27* mutation maps close to the centromere of chromosome 12 but, despite whole genomes sequencing approaches, the *PF27* gene product remained unknown (Kathir *et al*. 2003; Alford *et al*. 2013).

Taking a candidate approach, we searched the region near the *pf27* locus for genes with a possible flagella-related function and identified *ARMC2*, encoding an armadillo repeat protein conserved in organisms with motile cilia (Merchant *et al*. 2007). The mammalian homologue of ARMC2 has been linked to reduced lung function and male infertility but the precise role of ARMC2 in the assembly of motile cilia remains unknown (Soler Artigas *et al*. 2011; Coutton *et al*. 2019; Pereira *et al*. 2019). A novel *Chlamydomonas armc2* mutant shares the RS phenotype of *p27* and expression of *Chlamydomonas* ARMC2 restored wild-type motility and the presence of radial spokes in both *armc2* and *pf27* flagella, revealing that *PF27* encodes ARMC2. Fluorescent protein (FP)-tagged ARMC2 and the RS subunit RSP3 co-migrate on anterograde IFT trains in regenerating *Chlamydomonas* flagella whereas IFT of RSP3 was abolished in *armc2*. We conclude that ARMC2 is an adapter linking RSs to IFT to ensure their transport into flagella. Thus, IFT of ODAs, IDAs I1/f and RSs, three large axonemal substructures pre-assembled in the cell body, requires adapters with single cargo-specificity. As these adapters enter flagella and bind to IFT in the absence of their respective cargoes, we propose that the regulation of IFT-adapter interaction is a critical step for controlling cargo import into flagella by IFT.

## Results

### PF27 *encodes ARMC2*

The *pf27* locus was mapped to the midpoint between the *ODA9* and *TUB2* locus near the centromeric region of chromosome 12, placing it in the vicinity of the Cre12.g559250 gene, which encodes a 14-3-3 protein (Kathir *et al*. 2003). In the Phytozome genome browser (https://phytozome.jgi.doe.gov/pz/portal.html#), we inspected this region for genes with a potential role in flagella and identified Cre12.g559300 as a potential candidate. Cre12.g559300 encodes the armadillo-repeat protein ARMC2 (annotated as ARM1 in Phytozome), which is conserved in organisms with motile cilia (Li *et al*. 2004; Merchant *et al*. 2007). From the CLiP library we obtained the mutant strains LMJ.RY0402.155726 and LMJ.RY0402.083979, which have insertions in the 11^th^ and last intron of Cre12.g559300, respectively (Fig. 1A) (Li *et al*. 2019). Strain LMJ.RY0402.155726 had paralyzed flagella displaying residual jerky movements resembling *pf27* and we refer to this strain as *armc2*; the other strain swam normally and was not analyzed further (Fig. 1B, movie 1). Western blot analysis showed reduced levels of the radial spoke proteins RSP3 and nucleoside diphosphate kinase 5 (NDK5 aka RSP23) in flagella of *pf27* and *armc2* (Fig. 1C, D, Fig. 1 - Supplement 1A). In comparison to wild-type flagella, the slower migrating phosphorylated forms of these proteins, which can be separated by long runs on 6% acrylamide gels, were less abundant in *armc2* flagella, as previously described for *pf27* (Fig. 1 - Supplement 1A) (Huang *et al*. 1981). To visualize the distribution of radial spokes in *armc2* flagella, we generated an *armc2 pf14* RSP3-NG strain by genetic crosses. *PF14* encodes the RS protein RSP3, which is critical for radial spoke assembly and transport into flagella (Diener *et al*. 1993; Lechtreck *et al*. 2018). In control cells, RSP3-NG is present essentially along the length of flagella (Fig. 1 - Supplement 1B) (Lechtreck *et al*. 2018). In contrast, RSP3-NG was concentrated in the proximal region of the *armc2* flagella; it then tapered off and was largely absent from a large distal segment of flagella, similar to observations in *pf27* (Figs. 1E a – f, F d-f, S1B; (Alford *et al*. 2013)). In conclusion, *armc2* and *pf27* flagella share the same RS-related biochemical, structural and functional defects.

**Figure 1).**
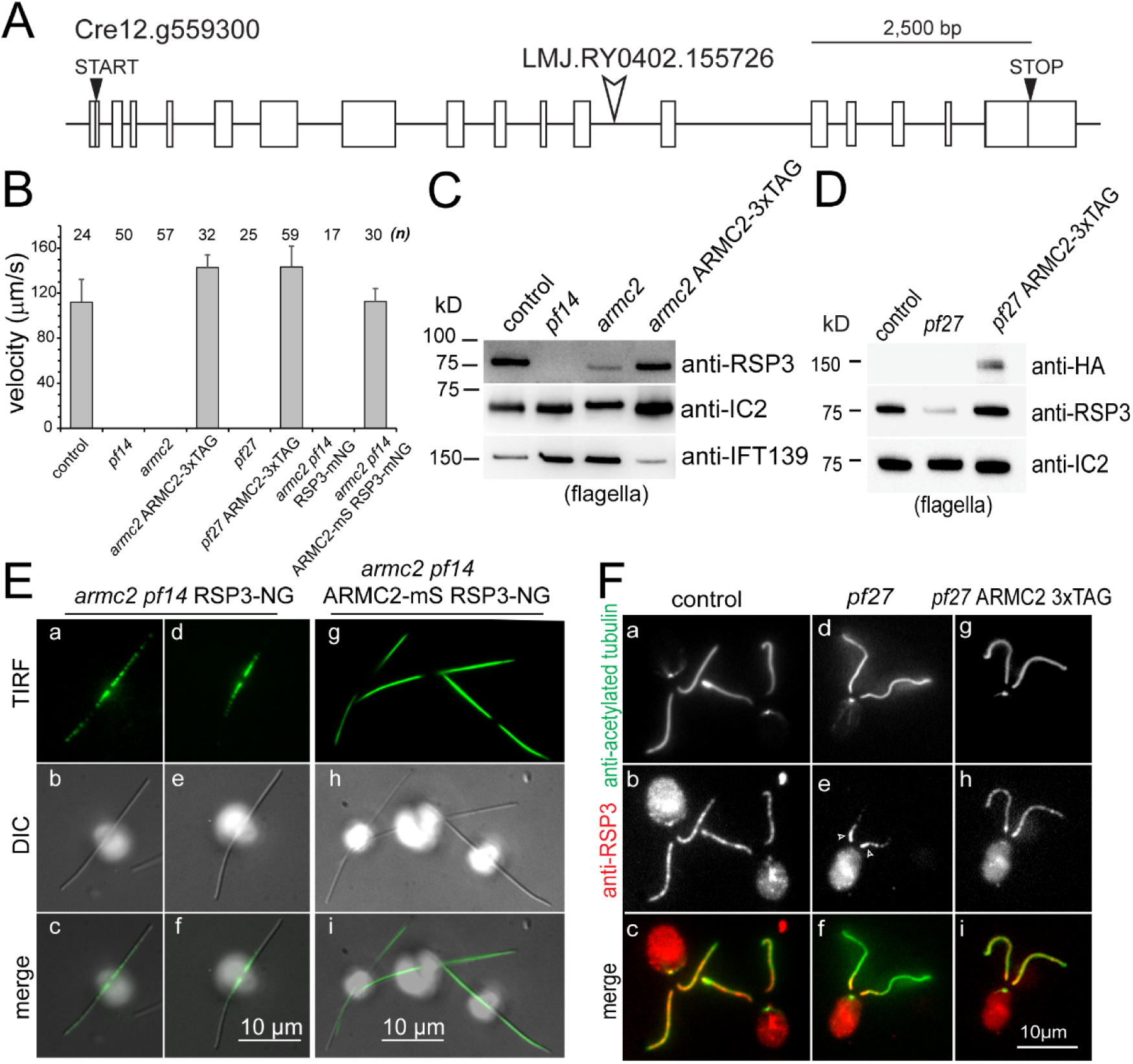
PF27 encodes ARMC2. A) Map of the *ARMC2* gene. The open arrowhead indicates the position of the insertion in the CLiP mutant LMJ.RY0402.155726. B) Average swimming velocity of the strains as indicated. The standard deviation and the number of cells analyzed are indicated. C) Western blot analysis of isolated flagella of control, the RSP3 mutant *pf14, armc2*, and the *armc2* ARMC2-3xTAG rescue strain with antibodies to RSP3 and as loading controls, the outer arm dynein subunit IC2 and IFT139. Note accumulation of IFT139 in *pf14* and *armc2* as previously reported for paralyzed central pair mutants of *Chlamydomonas*. D) Western blot analysis of isolated flagella of control, *pf27*, and the *pf27* ARMC2-3xTAG rescue strain with antibodies to RSP3, anti-HA and anti-IC2, as a loading control. Anti-HA was used to document expression of ARMC2-3xTAG (the 3xTAG encompasses a triple HA tag). A representative Western blot of two biological replicates is shown. E) DIC and TIRF imaging of live cells showing the distribution of RSP3-NG in the *armc2 pf14* mutant background (a – f) and a derived strain rescued by expression of ARMC2-mS (g-i). Bars = 10 μm. F) Immunofluorescence staining of methanol-fixed control (a – c), *pf27* (d – f) and *pf27* ARMC2-3xTAG (g – i) cells stained with anti-acetylated-α-tubulin (a, d, g) to visualize flagella and affinity-purified anti-RSP3 (b, e, h); merged images are shown in the bottom row (c, f, i). Arrows in e, residual RSP3 near the proximal end of the *pf27* flagella. Bar = 10 μm.

Using PCR, we engineered an *ARMC2* genomic construct encompassing the 11.3 kb *ARMC2* gene, approximately 1kb of each the 5’ and the 3’ flanking regions, and the aph7” selectable marker gene conferring resistance to hygromycin (Fig. 1 - Supplement 1C). An NG-3xHA-6xHis tag, here referred to simply as 3xTAG, or an mScarlet (mS) tag were added upstream of the *ARMC2* Stop codon (Fig. 1 - Supplement 1C). Transformation of *armc2* with the ARMC2-3xTAG plasmid restored wild-type motility and Western blotting showed near wild-type levels of RSP3 in the flagella (Fig. 1 B, C). TIRF microscopy showed that the normal distribution of RSP3-NG along the length of flagella was reestablished in the *armc2 pf14* ARMC2-mS RSP3-NG rescue strain (Figs. 1E g – i). Importantly, expression of ARMC2-3xTAG in *pf27* restored the presence and distribution of RSP3 in flagella and *pf27* ARMC2-3xTAG cells swam with wild-type motility (Figs. 1B, D, F g-i). Thus, introduction of the *ARMC2* gene rescues the RS defects in both *pf27* and *armc2* indicating both mutants are allelic and that mutations in *ARMC2* underly the *pf27* phenotype.

### ARMC2/PF27 is a conserved 107-kD armadillo-repeat protein

*Chlamydomonas ARMC2* is predicted to encode a 107 kD protein. We used mass spectrometry of ARMC2-3xTAG affinity-purified from the *armc2* ARMC2-3xTAG strain to confirm the coding sequence of ARMC2. The purified fusion protein migrated at approximately 160 kD in Western blots, consistent with its predicted size (107 kD and 34 kD for the tag; Fig. 1 - Supplement 1D). Using Ni-NTA-purification from whole cell extracts and anti-NG-nanobody trap purification from flagellar extracts, we identified a total of 17 unique ARMC2 peptides. Together with additional peptides obtained by mining other proteomic studies (Zhao *et al*. 2019; Picariello *et al*. 2020), the experimental peptides covered 42% of the predicted protein and were distributed through all but 1 (i.e., the 14^th^) of the 17 predicted ARMC2 exons (Fig. 1 - Supplement 1E).

*Chlamydomonas* and human ARMC2 are reciprocal best hits in protein BLAST searches with an E value of 4E-50. The N-terminal ∼400 residues of ARMC2 (∼300 residues of the smaller human Armc2 isoform CRA_b) are predicted to be mostly intrinsically disordered by IUPred2A (IUPred2A (elte.hu)) and include the ten phospho-sites identified by phosphoproteomics (Fig. 1 - Supplement 1E, F) (Wang *et al*. 2014; Erdos and Dosztanyi 2020). The C-terminal ∼600 residues of ARMC2 are predicted to be largely α-helical and encompass three armadillo repeats. Similar to previous efforts using whole genome sequencing, we failed to identify the causal genetic defect in *pf27* by sequencing of genomic PCR products; a possible contributing factor is the highly repetitive nature of the centromeric DNA.

### ARMC2 is highly enriched in regenerating flagella

The *pf27* and *armc2* mutants, radial spokes assembly onto the axoneme is rather incomplete but for the proximal region of the flagella (Alford *et al*. 2013). To address the question how ARMC2/PF27 promotes radial spoke assembly along the length of flagella, we turned to TIRF imaging of ARMC2-3xTAG expressed in the *armc2* mutant. In full-length flagella, we observed only few ARMC2-3xTAG particles moving by diffusion and, occasionally, by anterograde IFT (0.9 IFT events/minute/flagellum, SD 1.6 events/minute/flagellum, n=16; Figs. 2A a, B). In contrast, anterograde IFT of ARMC2-3xTAG was frequent in regenerating flagella with an average transport frequency of 44 events/minute/flagellum (SD 16.2 events/minute/flagellum, n=34) approaching those observed for GFP-tagged IFT itself (∼60-80/min; Fig. 2Ab, B) (Kozminski *et al*. 1993; Reck *et al*. 2016; Wingfield *et al*. 2017). Typically, ARMC2-3xTAG moved in one processive run from the flagellar base to the tip by anterograde IFT (green arrows in Fig. 2C a-c). As previously described for IFT proteins and other cargoes, ARMC2-3xTAG dwelled at the flagellar tip on average for 2.3 s (SD 1.8 s, n = 67; white brackets in Fig. 2C a - c) (Wren *et al*. 2013b; Chien *et al*. 2017). Dwelling and high-frequency transport resulted in the formation of a pool of ARMC2-3xTAG at the tip of regenerating flagella, but not full-length (Fig. 2 - Supplement 1A). Once released from IFT, ARMC2-3xTAG diffused swiftly into the flagellar shaft (white arrowheads in Fig. 2C a – c; Movie 2). Retrograde IFT of ARMC2-3xTAG was rare (Fig. 2B, red arrow in Fig. 2C d) indicating that the protein returns mostly by diffusion to the cell body as previously described for the anterograde motor kinesin-2 and IDA3, the cargo adapter for inner dynein arm I1/f (Chien *et al*. 2017; Hunter *et al*. 2018). Typically, ARMC2-3xTAG bleached in a single step indicating that only a single copy of the tagged protein was present on an individual IFT train (dashed circle in Fig. 2C b). However, trains carrying two copies of ARMC2-3xTAG were also observed (Fig. 2C e). Similar to other proteins transported by IFT, ARMC2-3xTAG displayed a variety of less frequent behaviors such as unloading from anterograde trains along the length of the flagella and re-binding to subsequent trains (not shown).

**Figure 2).**
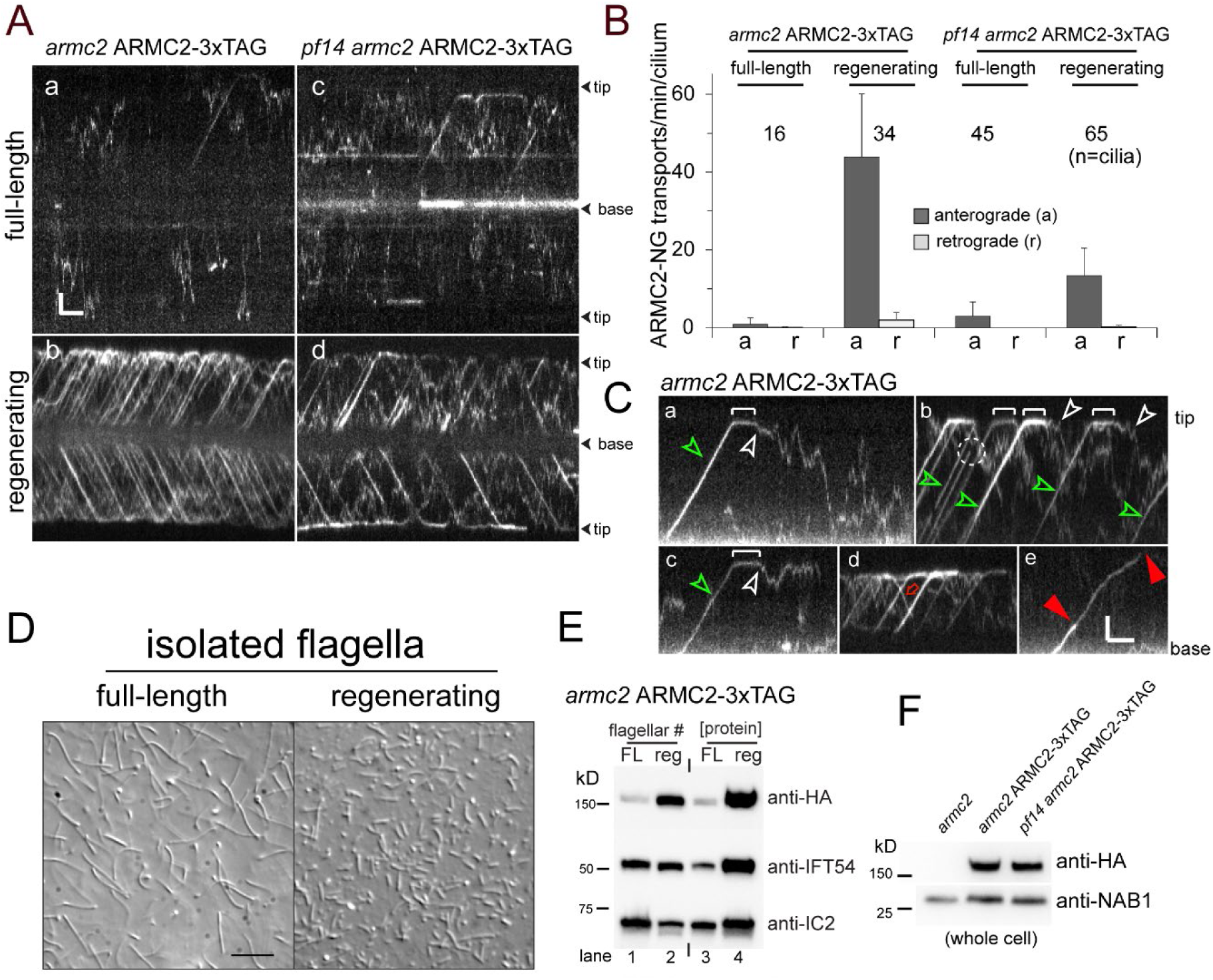
ARMC2-3xTAG is highly enriched in regenerating flagella. A) TIRF imaging of ARMC2-3xTAG in the *armc2* background (a, b) and the *pf14 armc2* double mutant background (c, d) in full-length flagella (a, c) and in regenerating flagella (b, d). Bars = 2 s 2 μm. The flagellar tips and bases are indicated. B) Bar graph showing the average frequencies of anterograde and retrograde transport of ARMC2-3xTAG in full-length and regenerating flagella of the *armc2* ARMC2-3XTAG and the *pf14 armc2* ARMC2-3xTAG strain. The standard deviation and the number of flagella analyzed are indicated. C) Kymograms of ARMC2-3xTAG in late regenerating flagella. The white brackets in a – c mark the dwell time of individual ARMC2-3xTAG particles between arrival at the tip by anterograde IFT and the onset of diffusion (white arrowheads). Green arrowheads in a - c, anterograde transport of ARMC2-3xTAG, red open arrow in d, retrograde IFT of ARMC2-3xTAG; red arrowheads in e, stepwise bleaching of ARMC2-3xTAG indicating for the presence of 2 copies. In c, a single step bleaching event is marked by a dashed circle. Bars = 2 s and 2 μm. D) DIC images of full-length and regenerating flagella of the *armc2* ARMC2-3xTAG strain. Regenerating flagella were harvested approximately 22 minutes after the deflagellation by a pH shock. Bar = 10 μm. E) Western blot analysis of the full-length and regenerating flagella shown in C with the antibodies indicated. On the left side, an equal number of flagella were loaded and on the right side, approximately equal loading of protein was attempted. F) Western blot comparing the presence of ARMC2-3xTAG in the *armc2* ARMC2-3xTAG and the *pf14 armc2* ARMC2-3xTAG strain. Antibodies to the cell body protein Nucleic Acid Binding protein 1 (NAB1) were used as a loading control.

To determine the degree of ARMC2-3xTAG accumulation in growing flagella, we isolated full-length and regenerating flagella from the *armc2* ARMC2-3xTAG strain for Western blot analysis (Fig. 2D, E). When a similar number of flagella, i.e., one regenerating for each full-length flagellum, were loaded, ARMC2-3xTAG was enriched about 14x in the regenerating flagella (Fig. 2E, lane 1 and 2). In contrast, the IFT particle protein IFT54 was similarly abundant in both samples, in agreement with previous observations that the amount of IFT proteins in flagella is largely independent of flagellar length (Marshall *et al*. 2005). When approximately similar amounts of protein were loaded, i.e., several short flagella for each full-length flagellum, both IFT54 and especially tagged ARMC2 were enriched in regenerating flagella (Fig. 2E, lane 3 and 4). Thus, ARMC2 is highly enriched in growing flagella.

To test if IFT of ARMC2 depends on the presence of intact RSs, we imaged ARMC2-3xTAG in a *pf14 armc2* double mutant. ARMC2-3xTAG moved by IFT in *pf14* flagella and the frequency of its transport was upregulated in regenerating *pf14* flagella (Fig. 2A d – f, B). In conclusion, IFT of ARMC2-3xTAG and its regulation by flagellar length do not require the presence of RSP3. However, IFT of ARMC2-3xTAG during flagellar regeneration in the *pf14* background was less frequent than in cells with intact RSs. Western blot analysis of whole cell samples showed similar amounts of ARMC2-3xTAG in the *armc2* ARMC2-3xTAG rescue strain and the *pf14 armc2* ARMC2-3xTAG strain (Fig. 2F). Thus, the presence of intact RSs could promote IFT of ARMC2-3xTAG. Similar to other strains with paralyzed flagella and large structural defects of the axoneme, *pf14* flagella regenerate slower than wild-type strains and Western blotting showed an accumulation of IFT proteins in full-length flagella (Figs. 1C, FIG. 2 - SUPPLEMENT 1B and not shown) (Lechtreck *et al*. 2013). Therefore, reduced IFT of ARMC2-3xTAG in *pf14* could also result from a more general imbalance of IFT rather than from the absence of intact spokes.

### RSP3-NG comigrates with ARMC2-mS during anterograde IFT

In growing flagella, ARMC2-3xTAG moves on anterograde IFT trains suggesting that it could assist in IFT of RSs, which are also transported more frequently during flagellar growth (Fig. 3A, B) (Lechtreck *et al*. 2018). 2-color TIRF imaging of the *armc2 pf14* ARMC2-mS RSP3-NG in the double-mutant-double-rescue strain revealed co-migration of the two tagged proteins during anterograde IFT (Fig. 3A, movie 3). In detail, 104 (80%) of 130 observed RSP3-NG anterograde transports observed over 1622 seconds in 20 regenerating flagella were accompanied by ARMC2-mS (Fig. 3A, C, Table 1). However, ∼82% of the ARMC2 transport were not accompanied by a RSP3-NG signal showing that ARMC2-mS transports were considerably more frequent than those of RSP3-NG in both full-length and regenerating flagella (Fig. 3A, C, Table 1). With respect to RSP3-NG transports lacking an ARMC2-mS signal, we note that in order to visualize IFT of RSP3-NG, we first had to photobleach RSP3-NG already incorporated into the axoneme. As mS is less photostable than NG, it is likely that some ARMC2-mS was photobleached during this preparatory step probably explaining the occurrence of some RSP3-NG transports without a co-migrating ARMC2-mS signal. To conclude, the transport frequency of ARMC2-mS exceeds that of its cargo RSP3-NG.

**Figure 3).**
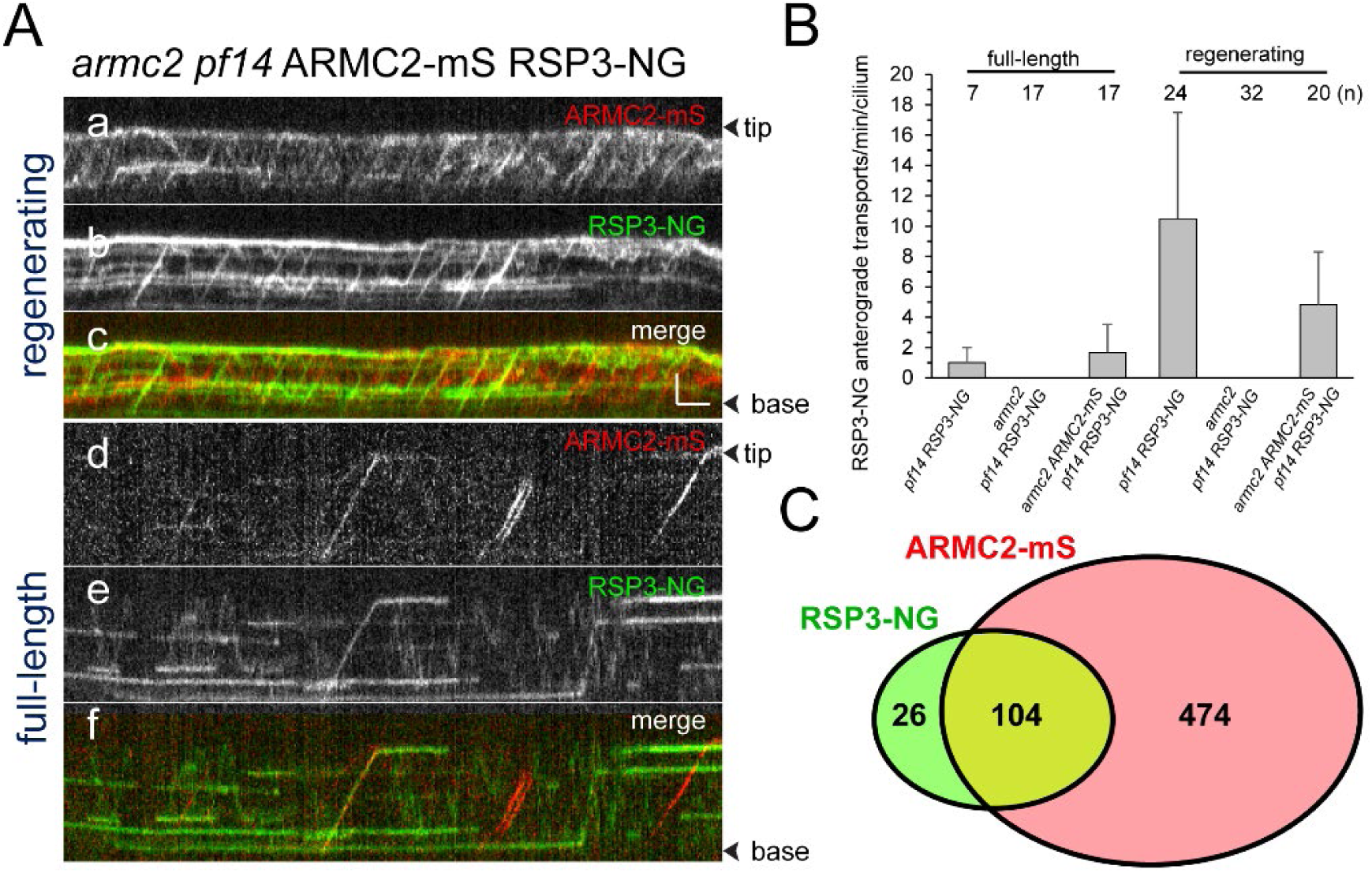
IFT of RSP3 requires ARMC2. A) Two-color TIRF imaging of a regenerating (a – c) and a full-length (d - f) flagellum of the *armc2 pf14* ARMC2-mS RSP3-NG strain. Horizontal trajectories result from residuals unbleached RSP3-NG in the axoneme. Bars = 2 s 2 μm. B) Bar graph showing the frequencies of RSP3-NG transports by anterograde IFT in full-length and regenerating flagella of the *pf14* RSP3-NG, *armc2 pf14* RSP3-NG and *armc2 pf14* ARMC2-mS RSP3-NG strains. The standard deviation and the number of flagella analyzed are indicated. C) Venn diagram showing the distribution of anterograde ARMC2-mS and RSP3-NG transports; the overlap area represents the co-transports corresponding to 82% of all RSP3-NG and 18% of the ARMC2-mS transports.

To analyze the behavior of the ARMC2-mS RSP3-NG complexes at the tip, we focused on the later stages of flagellar regeneration (>30 minutes after pH shock) when ARMC2-mS traffic was less dense increasing the chance of observing individual ARMC2-mS particles. Further, we photo-bleached the tip of the regenerating flagella in a subset of experiments to prevent unbleached RSP3-NG from accumulating as it incorporates into the elongating axoneme. The analysis of 15 such ARMC2-mS RSP3-NG complexes, showed that the release of ARMC2-mS from its dwell phase at the tip occurred concurrently with the onset of RSP3-NG movements (white arrowheads in Fig. 4A). While ARMC2-mS mostly diffused into the flagellar shaft, RSP3-NG moved to a somewhat more subdistal position, where it typically remained stationary for extended periods of time (12 of 15 events) potentially indicating stable docking to the axoneme. Diffusion of RSP3-NG deeper into the flagellar shaft (2 of 15 events) or return by retrograde IFT (1 of 15 events) were also observed (not shown). Thus, ARMC2 and RSP3 separate after arrival and dwelling at the tip with RSP3 remaining in the flagellum and ARMC2 returning to the cell body.

**Figure 4).**
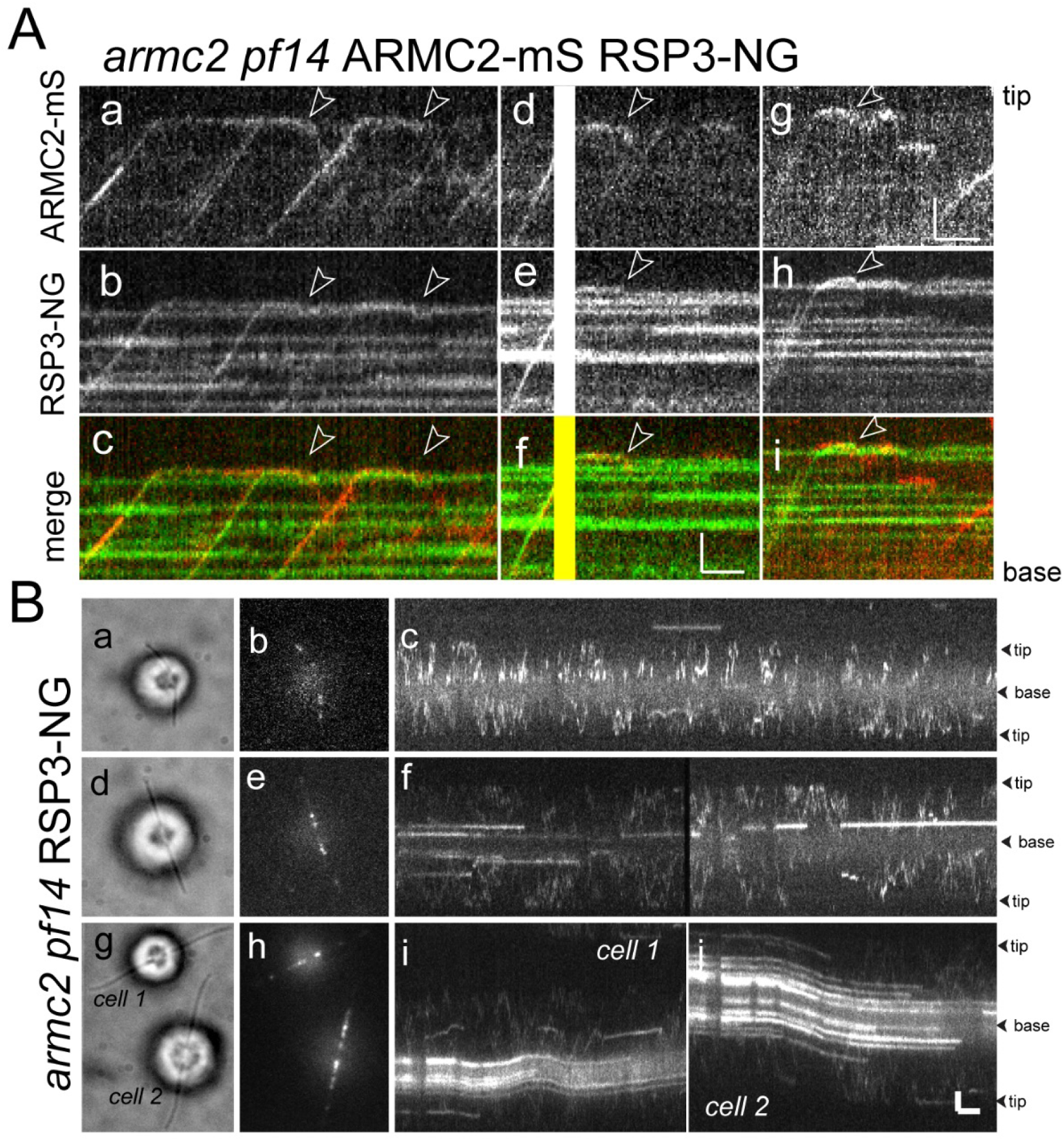
RSP3-NG ARMC2-mS complexes dissociate at the flagellar tip. A) Kymograms from simultaneous imaging of the cargo adapter ARMC2-mS (a, d, g) and its cargo RSP3-NG (b, e, h); the merged images are shown in c, f, and i. The end of the dwell phase and concomitant onset of ARMC2-mS and RSP3-NG movements are marked with white arrowheads. The white/yellow frames in d-f result from overexposure due to the use of the bleaching laser pointed at the other flagellar tip of the cell. Bars = 2 s and 2 μm. B) Brightfield (a, d, g) and TIRF (b, e, h) still images and corresponding kymograms (c, f, i and j) of early (a-c), mid (g-f) and late stage (g - j) regenerating *armc2 pf14* RSP3-NG cells. Bars = 2 s and 2 μm.

### IFT of RSP3-NG is abolished in the armc2/pf27 mutant

To test whether RSP3-NG moves by IFT in cells lacking ARMC2, we expressed RSP3-NG in an *armc2 pf14* double mutant and analyzed full-length and regenerating flagella (Fig. 4B). Some RSP3-NG particles moved inside flagella by diffusion and, as flagella elongated, the amount of RSP3-NG anchored in the proximal flagellar region increased (Fig. 4D)(Alford *et al*. 2013). But for rare ambiguous events, transport of RSP3-NG by IFT was not observed in the *armc2* mutant background. We conclude that ARMC2 is required for IFT of radial spokes (Fig. 4D).

For reasons unknown, motile cells occasionally emerged in *armc2* and *pf27* cultures. Over a few days, motile cells in such cultures progressed from slow swimming with RSP3-NG/-dTomato scattered along the flagella to fully motile with a near normal complement of tagged RSP3. For *armc2 pf14* RSP3-NG, such motile cells were still resistant to paromomycin indicating the continued presence of the insertion cassette in the *ARMC2* gene. In regenerating flagella of such cells, IFT of RSP3-NG was occasionally observed by TIRFM (not shown). The phenomenon was not further explored in this study.

### The IFT frequency of ARMC2 is regulated by flagellar length

The frequency of ARMC2-FP transports by IFT is upregulated in short growing flagella. To gain insights into how the frequency of ARMC2 transport is regulated, we generated long-zero cells by removing just one of the two flagella of a given cell by mechanical shearing. Such long-zero cells will regrow the missing flagellum while shortening the remaining one until both flagella are of approximately the same length (Rosenbaum *et al*. 1969; Ludington *et al*. 2012). Then, both flagella will regrow to full length. In all 19 *armc2* ARMC2-3xTAG long-short cells analyzed, the IFT frequency of ARMC2-3xTAG in the shorter flagellum exceeded that of the longer flagellum (on average 29.1 events/minute, SD 16 events/minute for the shorter vs. 8.9 events/minutes, SD 6.8 events/minutes for the longer flagellum; Fig. 5A, B). As the length difference between the short and long flagellum decreased, the difference in ARMC2-3xTAG transport frequency between the two flagella also diminished (Fig. 5A c and d, B). A similar behavior, i.e., flagellum-autonomous regulation of transport frequency by flagellar length, was previously documented for GFP-tubulin, DRC4-GFP, and the cargo adapter IDA3 (Craft *et al*. 2015; Hunter *et al*. 2018).

**Figure 5).**
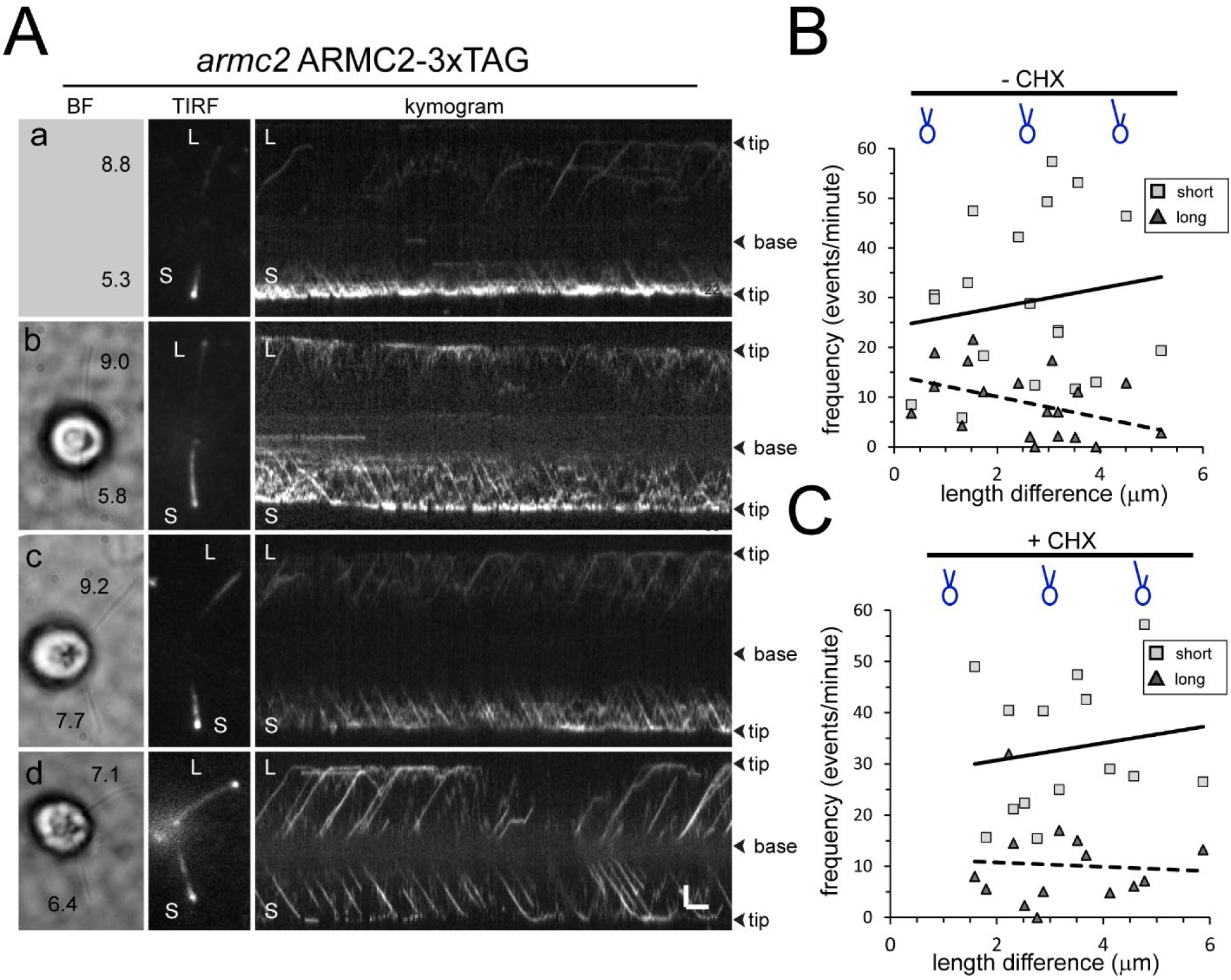
ARMC2-3xTAG transport is upregulated in short flagella. A) Gallery of brightfield (BF) and TIRF still images and the corresponding kymograms of long-short cells. No brightfield image was recorded for the cell shown in a. The length of the long (L) and short (S) flagella is indicated (in μm in a - d). Bars = 2 s and 2 μm. B) Plot of the ARMC2-3xTAG transport frequency in the short (squares) and the long (triangles) flagella against the length difference between the two flagella. Trendlines, solid for the long and dashed for the short flagella, were added in Excel. C) as B, but for cells treated with cycloheximide prior and during the experiment.

To test if the upregulation of ARMC2 transport during flagellar regeneration depends on *de novo* protein synthesis, we incubated cells in the protein synthesis inhibitor cycloheximide (CHX; 10 μg/ml) for 1 hour prior to generating long-short cells followed by regeneration in CHX-containing medium (Fig. 5C). The frequency of anterograde ARMC2-3xTAG transport in the shorter flagellum exceeded that of the longer flagellum of the same cell in all 14 long-short cells analyzed with average transport frequencies of 33 events/minute (SD 13.2 events/minute, n = 14) and 10.2 events/minute (SD 8.1 events/minute, n = 14) for short and long flagella, respectively (Fig. 5C, movie 4). Thus, ARMC2-3xTAG anterograde transport in sheared regenerating short flagella is approximately 4X more frequent than in the longer flagella of the same cell regardless of the cycloheximide treatment.

We also deflagellated cells by a pH shock in the presence of CHX and allowed them to regrow flagella in the presence of 10 μg/ml CHX for 90 minutes, at which point the flagella are ∼half-length and elongation had ceased due to the absence of protein synthesis (Rosenbaum *et al*. 1969). In such cells, the frequency of ARMC2-3xTAG transport was 14.4 events/minute (SD 4.3, n = 10) well above the frequency determined for full-length flagella (see Fig. 2B). Thus, high frequency ARMC2-3xTAG transport is indeed triggered by the insufficient length of the flagella rather than active growth of flagella. Taken together, the data indicate that cells possess a pool of ARMC2-3xTAG and that ARMC2-3xTAG will preferably attach to IFT trains that enter short flagella.

### The ARMC2 cell body pool

The distribution and dynamics of cargo adapters and cargo proteins in the cell body remains largely unknown. The data above established the presence of an ARMC2-FP pool in the cell body. To determine its distribution, we analyzed *pf14 armc2* RSP3-NG ARMC2-mS cells using epifluorescence, which revealed the presence of an ARMC2-mS pool near the flagellar base of most cells (Fig. 5 - Supplement 1A). To analyze the dynamics of the ARMC2 pool, we increased the incidence angle of the TIRF laser, allowing us to image ARMC2-3xTAG positioned deeper in the cell body (Fig. 5 - Supplement 1B). The ARMC2-3xTAG pool typically consisted of two closely spaced dots and we use a focused laser beam to photobleach one of the two dots (Fig. 5 - Supplement 1B and C). Due to the closeness of the two dots, the signal of the second dot was diminished during the bleaching step but could still be used as a control to estimate signal recovery. Analysis of a small number of *armc2* ARMC2-3xTAG and *pf14 armc2* ARMC2-3xTAG cells (n=3) showed that recovery of the ARMC2-3xTAG was incomplete and plateaued after approximately 20s suggesting a slow exchange rate of the bleached protein in the pool.

ARMC2-3xTAG cells with regenerating flagella were used to determine the dwell time of ARMC2 in the basal body-associated pool. After bleaching of the entire basal body-associated pool, IFT of ARMC2-3xTAG was interrupted for approximately 19 s (SD 7.7 s, n = 12, red bracket in Fig. 5 - Supplement 1E, top panel). A similar average length of the gap was observed in the *pf14 armc2* ARMC2-3xTAG (20 s, SD 10.7 s, n = 18; Fig. 5 - Supplement 1E, bottom panel) suggesting that ARMC2 dynamics in the pool, i.e., the rate by which it travels through the pool and attaches to IFT, are not altered by the absence of its cargo. After the gap, ARMC2-3xTAG traffic recommenced albeit at a reduced frequency in agreement with the incomplete recovery of the signal shown in Fig. 5 - Supplement 1D. Considerably shorter gaps were observed for several IFT proteins (∼ 6 s or less) and tubulin-GFP in similar bleaching experiments, (∼2 s) (Wingfield *et al*. 2017). Thus, ARMC2-3xTAG dwells in the basal body-associated pool considerably longer than the IFT proteins of the train, which will carry it into the cilium, suggesting that IFT proteins and ARMC2 are recruited through different routes.

### ARMC2, IDA3 and IC2 are stochastically distributed onto IFT trains

Similar to previous observations on the I1/f transport adapter IDA3 and several axonemal proteins (Wren *et al*. 2013a; Craft *et al*. 2015; Hunter *et al*. 2018), IFT of ARMC2-3xTAG is frequently observed in short flagella but progressively decreases as flagella approach full length. How axonemal proteins and their adapters are distributed onto the IFT trains remains unknown as the currently available cryo-EM structures of IFT trains were obtained by image averaging, which will cancel-out signals from substoichiometric train components such as cargoes (Jordan *et al*. 2018). One possible model is that a subset of trains is specialized to carry such axonemal cargoes, e.g., because they are in an “open” state allowing cargoes to adhere. Such “axonemal cargo trains” could frequently enter short flagella but only sporadically move into full-length flagella explaining the flagellar length-dependent decline in cargo IFT. The model predicts that different axonemal proteins preferentially travel on this subclass of trains. Alternatively, all trains are equally capable of binding cargo and the proteins will stochastically distribute onto the trains (Fig. 6A).

**Figure 6).**
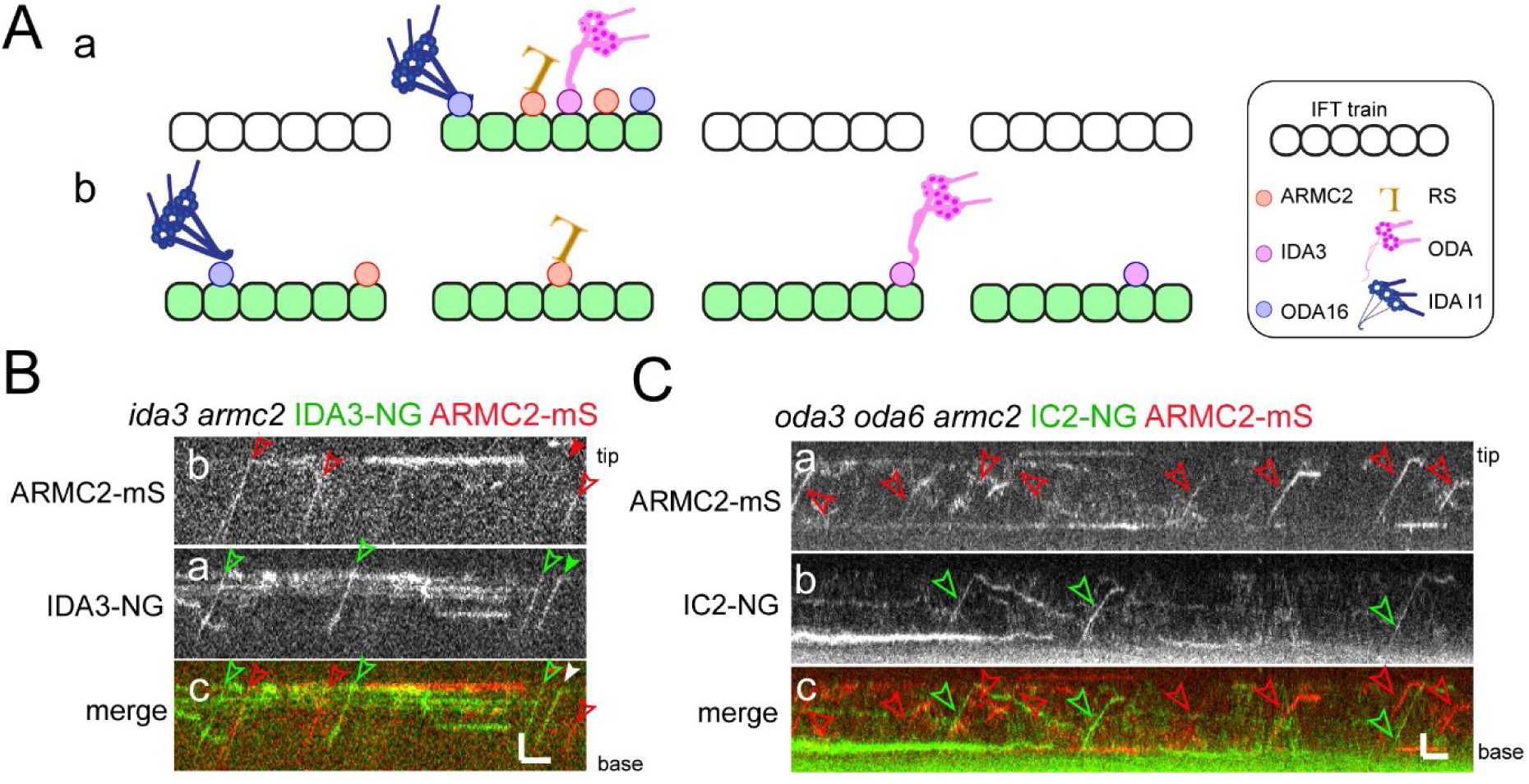
ARMC2-mS is transported independently of IDA3-NG and IC2-NG. A) Schematic presentation of two models for IFT-cargo interaction using RSs, ODAs and I1 IDAs and their adapters as examples. a) Most cargoes use a specific subset of IFT trains, which have a high propensity to bind axonemal proteins, for example, because they are in a hypothetical open configuration. b) All IFT trains are similarly capable of binding axonemal cargoes; thus, cargoes are stochastically distributed onto the trains. B) Kymograms of two-color TIRF imaging of ARMC2-mS and IDA3-NG in an *ida3 armc2* mutant cell. ARMC2-mS trajectories are marked with open red arrowheads and IDA3-NG trajectories with open green arrowheads. Filled arrowheads indicate a co-transport. Bars = 2 s and 2 μm. C) Kymograms of two-color TIRF imaging using the *oda3 oda6 armc2* IC2-NG ARMC2-mS strain. Trajectories of ARMC2-mS and IC2-NG transports are marked with red and green arrowheads, respectively. Bars = 2 s and 2 μm.

We generated strains expressing the adapter proteins ARMC2-mS and IDA3-NG in a corresponding *armc2 ida3* double mutant or expressing ARMC2-mS together with the essential outer dynein arm (ODA) subunit IC2-NG in an *oda3 armc2 oda6* strain (Fig. 6B, C). IC2-NG rescues the ODA-deficient *oda6* mutant and the ODA docking complex-deficient *oda3* mutant background was chosen because it interferes with the binding of ODAs to the axoneme preventing the accumulation of IC2-NG in flagella and thereby circumventing the need to photobleach flagella prior to imaging (Koutoulis *et al*. 1997; Dai *et al*. 2018). Live imaging of ODA16, the adapter required for OFA transport, has not yet been achieved. We focused on mid to late regenerating cells when the transport of these proteins was infrequent, reducing the probability that both proteins are present on a given train by chance. In 35 regenerating *armc2 ida3* ARMC2-mS IDA3-NG flagella analyzed over 905 seconds, we observed 243 ARMC2-mS and 106 IDA3-NG particles moving anterogradely by IFT, of which 20 were co-transports on one train (Table 1). For the *oda3 armc2 oda6* ARMC2-mS ODA6-NG strain, we analyzed 20 flagella for 1575 seconds and observed 82 ARMC2-mS and 78 IC2-NG particles, of which 3 were co-transported (Table 1). The lower frequency of ARMC2-mS transports in the latter experiment could reflect that more cells in later stages of regeneration were analyzed as indicated by the longer average length of cilia (8.3 μm ± 1.85 μm vs. 7.3 μm ± 1.5 μm). As anterograde IFT has a frequency of ∼60 trains/minute, the observed probability that a train caries both cargoes was 0.022 for ARMC2-mS (i.e., ∼2% of the trains carry AMC2-mS) and IDA3-NG and 0.0019 for ARMC2-mS and IC2-NG. These values are close to the calculated probabilities of 0.03 for ARMC2-mS/IDA3-NG and 0.0026 ARMC2-mS/IC2-NG by which such co-transport would occur by chance when the two cargoes are transported independently of each other without a preference for a subclass of IFT trains (see Table 1 and Materials and Methods for calculation). In contrast, RSP3-NG transport depends on ARMC2-mS and as expected ARMC2-mS and RSP3-NG were co-transported with a probability of 0.027 in full-length flagella significantly exceeding the value of 0.0023 calculated for stochastic co-transport of independent cargoes. In conclusion, ARMC2-mS, IDA3-NG and IC2-NG are transported independently of each other arguing against a specific subclass of IFT trains specialized to carry axonemal proteins but rather supporting a model in which axonemal cargoes are stochastically distributed onto the trains.

## Discussion

### ARMC2 is an adapter for IFT of RSs

IFT adapters could be defined as a proteins or protein complexes, which facilitate IFT of a cargo (complex) without being necessary for either IFT itself or the functionality of the cargo. Here, we show that IFT of RSs requires ARMC2 as an adapter. The notion is supported by the following observations: 1. In *pf27* and *armc2* mutants, the presence of RSs was limited to the most proximal region of flagella, which is likely explained by residual entry of RSs into flagella by diffusion, 2. IFT of tagged RSs was not observed in *armc2* mutants, and 3. tagged ARMC2 and the essential RS protein RSP3 typically co-migrated on anterograde IFT trains. In comparison to wild-type flagella, the phosphorylation of several spoke proteins is reduced in *pf27/armc2* (Huang *et al*. 1981). While details remain to be explored, it has been proposed that RS phosphorylation occurs near the flagellar tip (Gupta *et al*. 2012). Then, the altered phosphorylation of RS proteins in *pf27/armc2* could result from the RSs failing to reach the flagellar tip because they cannot attach to IFT. We conclude that ARMC2/PF27 ensures efficient delivery of RSs into *Chlamydomonas* flagella and to the flagellar tip by linking RSs to IFT trains.

Mutations in mammalian *Armc2* cause sperm malformations resulting in male infertility (Coutton *et al*. 2019). Immunofluorescence analysis of spermatozoa from the affected individuals revealed central pair defects as assessed by the loss of the central pair marker proteins SPAG6 and SPEF2. The spoke head protein RSPH1 was detected in the stunted mutant flagella but the presence of RSs was not further evaluated (Coutton *et al*. 2019). In mammalian cilia, radial spoke defects are often associated with a dislocation or even loss of the central pair apparatus (Antony *et al*. 2013; Kott *et al*. 2013). Thus, a deficiency of the RSs could also lead to central pair defects. So far, *Armc2* defects in human patients have been linked only to male infertility, suggesting that Armc2 is expendable for the assembly of motile cilia in the airways and ventricular system (Coutton *et al*. 2019). However, genome-wide interaction studies have linked *Armc2* variants to reduced lung function (Soler Artigas *et al*. 2011; Pereira *et al*. 2019) and Armc2 is highly expressed in ciliated epithelial cells (Uhlen *et al*. 2015). With a length of ∼7 μm, airway cilia are comparatively short and it seems possible that RSs enter them over time by diffusion in amounts sufficient to ensure some degree of motility. To conclude, further studies are required to determine if the role of *Chlamydomonas* ARMC2 in RS transport is conserved in other organisms.

### IFT of large axonemal complexes involves adapters

Just like the transport of RSs, IFT of ODAs and IDAs I1/f requires ODA16 and IDA3, respectively, as transport adapters. The flagella of *armc2/pf27, ida3* and *oda16* mutants specifically lack or have greatly reduced amounts of RSs, I1/f dynein or ODAs, respectively, indicating that these adapter proteins promote transport with single-cargo specificity. In conclusion, IFT of three major axonemal substructures, all multiprotein complexes preassembled in the cell body, involves specific adapters suggesting that the transport of other axonemal complexes, such as the other inner dynein arms, may also require adapter proteins.

The identification of a third cargo adapter involved in axonemal assembly allows us to compare their features: ODA16 and ARMC2 are well conserved in organisms with motile cilia and flagella while IDA3 is not; the latter, however, shows partial similarity to coiled-coil domain-containing protein 24 (CCDC24)(Hunter *et al*. 2018). All three proteins were not detected in the original proteomic analysis of fractionated *Chlamydomonas* flagella (Chlamydomonas Ciliary Proteins (chlamyfp.org)) indicating a low abundance in full-length flagella (Pazour *et al*. 2005). Similarly, EST coverage supporting the expression of the genes are very limited or absent (Albee *et al*. 2013). ODA16 is a WD-repeat protein with a small C-terminal intrinsically disordered region, ARMC2/PF27 encompasses armadillo repeats and has an extended N-terminal intrinsically disordered region, and IDA3 possesses several short coiled-coil regions interspersed in a largely intrinsically disordered protein (Taschner *et al*. 2017; Hunter *et al*. 2018). Intrinsically disordered regions often adopt a more defined structure upon binding to their partners (Dyson and Wright 2005). Thus, IFT adapters could fold once they interact with IFT and/or their cargoes. With the phosphorylated peptides mapped to its disordered region, phosphorylation in this region could hypothetically regulate ARMC2’s folding and molecular interactions (Johnson and Lewis 2001; Iakoucheva *et al*. 2004).

The interaction of ODA16 with the N-terminal region of IFT46 requires both its C-terminal intrinsically disordered region and the WD-repeats whereas the C-terminal region of ODA16 is expandable for ODA binding (Taschner *et al*. 2017). The IFT binding sites of ARMC2/PF27 and IDA3 for remain unknown. Potential candidates for IDA3 binding include the IFT-B proteins IFT56 and IFT57 as the flagellar levels of IDAs or IDA subunits are reduced in the corresponding mutants (Ishikawa *et al*. 2014; Jiang *et al*. 2017). Which subunits of the RSs, ODAs and IDAs interact with the respective adapter and thus mediate transport of the entire complex, remains to be explored. In *Drosophila*, RS genes are highly expressed in testis but only the *RSP3/PF14* orthologue *CG32392* is expressed in embryonic chordotonal neurons (ZUR LAGE *et al*. 2019). Interestingly, the transcript of *CG32668*, the fly orthologue of ARMC2, was abundant in chordotonal neurons but not detected in testis (Andrew Jarman, personal communication, August 2021). The development of 9+2 sperm flagella by an IFT-independent mechanism likely explains the absence of *Armc2* expression in testis. The expression of both *Armc2* and *CG32392* (*Rsp3*) in chordotonal neurons, which use the IFT pathway to assemble 9+0 cilia, could indicate that ARMC2 interacts with RSP3 and that RSP3 is part of these non-motile but mechanosensitive cilia, which possess inner and outer dynein arms. Similarly, some RS proteins are also present in 9+0 motile cilia (Sedykh *et al*. 2016).

### IFT cargo adapters provide an additional level to regulate cargo flux

ARMC2 and IDA3 are enriched in short growing flagella whereas only traces are present in full-length flagella, when cargo transport is contracted. ODA16 is present in full-length flagella (Ahmed and Mitchell 2005) and our analysis showed that is also enriched in growing flagella (Fig. 2 - Supplement 1C). IFT of ODA16 has not been observed directly but its interaction with IFT46 is well supported by genetic, biochemical, and structural data (Ahmed *et al*. 2008; Hou and Witman 2017; Taschner *et al*. 2017). Thus, it is reasonable to assume that ODA16 moves by IFT and moves more frequently during flagellar regeneration, just as IFT of ODAs is upregulated in growing flagella (Dai *et al*. 2018). ODA16 is present in *oda2* flagella, which lack ODAs (Ahmed and Mitchell 2005; Hunter *et al*. 2018), and IDA3 and ARMC2/PF27 continue to move by IFT in the absence of their respective cargoes (Hunter *et al*. 2018). Further, the transport frequencies of IDA3 and ARMC2/PF27 are still upregulated in short flagella of mutants lacking the respective cargo. In the *armc2 pf14* ARMC2-mS RSP3-NG double mutant double rescue strain, numerous ARMC2-mS solo transports were observed, in addition to ARMC2-mS RSP3-NG cotransports. Thus, even when RSs are available, ARMC2 is often loaded onto IFT without its cargo. In a hypothetical model, cells regulate ARMC2-IFT interactions and RSs, when available, will latch on to IFT using the binding sites generated by ARMC2. If correct, the frequency by which these axonemal building blocks are transported into flagella would be controlled to a substantial part by the regulation of IFT-adapter interactions. The model does not exclude that adapter-cargo interactions are regulated as well. Binding of the BBSome to IFT trains is regulated by three small GTPases, IFT22, IFT27 and RabL2 and BBSome-dependent export requires ubiquitination of its GPCR cargoes (Eguether *et al*. 2014; Desai *et al*. 2020; Shinde *et al*. 2020; Xue *et al*. 2020; Duan *et al*. 2021). Functional analysis of the phosphorylation sites in the unordered region of ARMC2 could provide insights into the regulation of IFT-ARMC2-RS interactions.

### Unrelated cargoes bind stochastically to IFT trains

Most IFT trains are densely loaded with axonemal cargoes and adapters during early flagellar growth (Wren *et al*. 2013a; Craft *et al*. 2015; Dai *et al*. 2018; Lechtreck *et al*. 2018). The transport frequencies decline as flagella approach full length and a given cargo is present on only a small subset of the anterograde trains. It is currently unknown whether axonemal cargoes are randomly distributed onto all available IFT trains or are preferably transported on a specialized subclass of “active” or “open” IFT trains, which are abundant during rapid flagellar growth but scarce during maintenance of full-length flagella. The latter model predicts that unrelated axonemal cargoes will frequently ride on the same IFT trains, which should be particularly apparent in late regenerating flagella when the transport frequencies of axonemal proteins are low. However, the adapters ARMC2 and IDA3 and the ODA subunit IC2 travel apparently independently of each other. If these proteins are representative, cargoes appear to be distributed stochastically onto IFT trains arguing against a subclass of cargo-carrying trains and against a regulation of cargo binding at the level of entire trains. The regulation of IFT-adapter-cargo interaction could instead occur at the level of individual IFT complexes or even independently of IFT trains altogether, for example, by modifications of the adapters and/or cargoes. Extrapolating our observations on ARMC2, IDA3 and the BBSome, we predict that the trains carry a random mix of cargo adapters and cargoes, which are substoichiometric to IFT itself and that this mix-and-match “IFT corona” will be prominent on trains entering short growing flagella but downsized in full-length flagella.

## Materials and methods

### Strains and culture conditions

The previously described *pf27* (CC-1387), *pf14* (CC-613) mutant strains and the CliP strains LMJ.RY0402.083979 and LMJ.RY0402.155726 (i.e., *armc2*) are available from the Chlamydomonas Resource Center (RRID:SCR_014960). The *pf14* RSP3-NG, IDA3-NG and the *oda3 oda6* IC2-NG strains were described previously (Zhu *et al*. 2017; Dai *et al*. 2018; Hunter *et al*. 2018). The *armc2 pf14* ARMC2-3xTAG, *armc2 pf14* ARMC2-mS RSP3-NG, *armc2 oda3 oda6* ARMC2-mS IC2-NG, and *ida3 armc2* IDA3-NG ARMC2-mS strains were generated by mating. Progeny with the desired combination of alleles were identified using a combination of selection (CliP mutants are resistant to paromomycin and the ARMC2 transgenes confer hygromycin resistance and fluorescence), PCR, fluorescent imaging, and western blotting. The *ida3* mutation generates a SfcI site and we used the primers ida3f (ATTTGGACGGAGCCTTGAC) and ida3r (TGTTTCGCACGCCTTCA) to amplify the genomic region flanking this site followed by restriction digest with Sfc1 to track the *ida3* mutant allele (Hunter *et al*. 2018)

Cells were maintained in M medium at a 14:10 hours light:dark cycle at 24 °C. The cells maintained in unaerated flask for imaging, transformation, and phenotypical analysis; cultures were aerated with air supplemented with 0.5% CO_2_ for flagellar isolation.

### ARMC2 cloning

The cloning scheme for the 13.6-kB ARMC2 genomic DNA was depicted in the supplemental data (Figure S1). Briefly, using PCR and purified genomic DNA as a template, five genomic DNA fragments of the ARMC2 gene were amplified. Fragment 1 of 4.6 kB, including 1 kB 5’ flanking sequence, was amplified with primer S1** and AS2 and cloned into the pGEM-T vector by AT cloning. Fragment 2 of 3.4 kB and fragment 3 of 3.1 kB were amplified using primer pairs S3*/AS3* and S4/AS4 HindIII, respectively, AT cloned and then combined into one plasmid by releasing fragment 3 with a XhoI/SacI digest and ligating it into the pGEM-T- fragment 2 plasmid digested with the same enzyme. This resulted in a plasmid with the 6.5 kB fragment 2’ segment. The HindIII restriction sequence in the primer in an intron was created for ligation with the downstream fragment. Similarly, fragments 4 and 5 were amplified using primer pairs HindS6.1/AS5Xho and XhoS7/AS6 and AT cloned into pGEM-T. An XhoI restriction sequence was added into the primers so that fragment 5, released by a XhoI and SacII digest, can be ligated with fragment 4 in the pGEM-T plasmid digested with the same enzymes. This resulted in the plasmid containing the 2.5 kB fragment 3’. The tag (3xTAG or mS) was inserted into the Xho1 site of fragment 3’ and the correct orientation was verified by restriction digest. Then, three fragments - including fragments 2’ and 3’, respectively released by Spe1 and HindIII digest and HindIII and NdeI digest, along with a 1.7 kB fragment conferring the hygromycin (Hyg)-resistance (Zhu *et al*. 2017), were inserted into the pGEMT-fragment 1 plasmid digested with SpeI and NdeI. The ligation mixture was transformed into DH10B competent *E. coli* cells (New England Biolab, MA). The final pGEM-T-ARMC2 construct was confirmed by restriction digest.

List of primers:

S1**- CCGCCTGCACCCTTATCGCTGCCTCTGTCCCTCTTCC

AS2 – CCTGTTCCGCACGCTGGTCTACCGTCTACC

S3*- CGAGGCGGTGAGCGAGCACGTGTTCCGACTCATG

AS3*- GCCTCACGGTACCGTGAGCACATGCATGGGTTTGC

S4 - CGCAACCCCCGCTACTCTAACCTCGAGG

AS4Hind- CAG<u>AAGCTT</u>GAAGCCCGAAAGCTGACGAAGTGGG

HindS6.1-GAG<u>AAGCTT</u>ACCTACCTGGGTCTTGACATGCCCTGTCC

AS5Xho- C<u>CTCGAG</u>CTCCGGCAACGCCTCCAGCTCC

XhoS7- C<u>CTCGAG</u>TAGGGGCCCTTGCTTAGGGAATTCAGGG

AS6- CTCGCTTTCACAACTCCAGGGTGCCCATGC

### Generations of ARMC2 transgenic strains

Purified plasmids of the ARMC2 genomic construct were transformed into *pf27* and *armc2* cells using the glass beads method (Kindle 1990; Zhu *et al*. 2017). Transformants were selected on plates containing 10 μg/ml of hygromycin and the resistant clones were suspended in 10 mM Hepes and further screened for motility and fluorescence.

### Flagellar isolation and Western blotting

For Western blot analyses, cells were washed in 10 mM Hepes, resuspended in 10 mM Hepes, 5 mM MgSO_4_, 4% sucrose (w/v) and deflagellated by the addition of dibucaine. After removing the cell bodies by two differential centrifugations, flagella were sedimented at 40,000 x g, 20 minutes, 4 °C as previously described (Witman 1986). To obtain short regenerating flagella, cells were deflagellated by a pH shock, transferred to fresh M medium, stored on ice for ∼15 minutes, diluted with M medium and allowed to regenerated flagella for ∼18 minutes in bright light with agitation. Then, cells were sedimented, washed once in 10 mM Hepes and deflagellated as described above. Flagella were dissolved in Laemmli SDS sample buffer, separated on Mini-Protean TGX gradient gels (BioRad), and transferred electrophoretically to PVDF membrane. After blocking, the membranes were incubated overnight in the primary antibodies; secondary antibodies were applied for 90-120 minutes at room temperature. After addition of the substrate (Femtoglow by Michigan Diagnostics or ECL Prime Western Blotting Detection Reagent by GE Healthcare), chemiluminescent signals were documented using A BioRad Chemi Doc imaging system. Western blots shown in Fig. 1 – Supplement 1 were documented using film and those in Fig. 2 – Supplement 1 – panel B using an Alpha Innotech FluorChem Q chemiluminescence imaging system. The following primary antibodies were used in this study: rabbit anti-RSP3 (1:800) and, for immunofluorescence, affinity-purified anti-RSP3 (1:100; (Williams *et al*. 1986)), rabbit anti-NDK5 (1:1,000; (Chung *et al*. 2017)), rabbit anti-IFT54 (1:800)(Wingfield *et al*. 2017), rabbit anti-NAB1 (1: 5,000; Agrisera), rabbit anti-GFP (A-11122; ThermoFischer), mouse monoclonal anti-IFT139 (1:400)(Cole *et al*. 1998), and mouse monoclonals anti-IC1 (1:xxx) and IC2(King and Witman 1990), rat monoclonal anti-HA (1:800, clone 3F10 Roche/Sigma).

### Indirect immunofluorescence

For indirect immunofluorescence, cells were sedimented, resuspended in HMEK, allows to settle onto polyethyleneimine (0.2%) coated multiwell slides for 1-2 minutes and submerged into - 20°C methanol for 8 minutes. The slides were air-dried, blocked (1% BSA in PBS-T), washed with PBS, incubated with the primary antibodies in blocking buffer overnight, washed, stained with secondary antibodies (1:800 AlexaFluor anti-rb-565 and anti-mo-488; Invitrogen), washed in PBS-T, submerged briefly in 80% ethanol, airdried and mounted in ProlongGold (Invitrogen).

For widefield epifluorescence microscopy, images were taken using a 60x 1.49 objective, an Eclipse Ti-U microscope (Nikon) equipped with a Lumen200 light source (PRIOR) and filters for FITC and TexasRed. Alternatively, images were taken using a 40 X Plan Fluor lens on an Eclipse E600 microscope (Nikon) equipped with a DFC9000 GT sCMOS camera and the accompanied imaging software (Leica, Wetzlar, Germany). The bandpass of filters was Ex: 460-500 nm, Dm: 505 nm, Em: 510-560 nm for NeonGreen (NG) and Ex: 543-568 nm, Dm: 570 nm, Em: 579-612 nm for mSc. The illuminator was a 4-channel SOLA SM 365 LED light engine with excitation peaks at 365, 470, 530, 590 nm (Lumencor, Beaverton, OR).

### Live cell microscopy

For TIRF imaging, we used an Eclipse Ti-U microscope (Nikon) equipped with 60× NA1.49 TIRF objective and through-the-objective TIRF illumination provided by a 40-mW 488-nm and a 75-mW 561-nm diode laser (Spectraphysics) as previously described (Lechtreck 2013; Lechtreck 2016). The excitation lasers were cleaned up with a Nikon GFP/mCherry TIRF filter cube and the emission was separated using an Image Splitting Device (Photometrics DualView2 with filter cube 11-EM) supplemented with an et595/33m filter to reduced chlorophyll autofluorescence and a meniscus lens to adjust for focus. Images were mostly at 10 fps using an iXON3 (Andor) and the NIS-Elements Advanced Research software (Nikon). Data in Fig. 6 were documented using μManager instead of NIS-Elements. The optical set-up for the focused laser beam used for FARP analysis has been previously described set-up by Wingfield et al. (2017). FIJI (National Institutes of Health) was used to generate kymograms using either the build-in Multi Kymogram tool or the KymoResliceWide plugin (https://imagej.net/KymoResliceWide). The Plot Profile tool was used to analyze signal intensity and Microsoft Excel was used for statistical analysis. Adobe Photoshop was used to adjust image contrast and brightness, and figures were prepared in Adobe Illustrator.

Observation chambers for *C. reinhardtii* were constructed by applying a ring of vacuum grease or petroleum jelly to a 24 ° 60 mm No. 1.5 coverslip; 10 μl of cell suspension were applied and allowed to settle for ∼1 minute. Then, the chamber was closed by inverting a 22 × 22 mm no. 1.5 cover glass with ∼5-10 μl of 5 mM HEPES, pH 7.3 supplemented with 3-5 mM EGTA onto the larger coverslip. Cells were imaged through the large cover glass at room temperature. Regenerating cells were obtained as follows: Cells in fresh M-medium were deflagellated by a pH shock, sedimented, resuspended in a small volume of M medium and stored on ice for 15 minutes or until needed. Then, cells were diluted with M medium and agitated in bright light at room temperature. Aliquots were analyzed by TIRF at various time points. To generate long-short cells, cells in M medium were chilled on ice for ∼3 minutes and then passed repeatedly (5-8 x) through a 26G½ needle using a 1-ml syringe (Ludington *et al*. 2012). The presence of spinning long-zero cells was verified by microscopy and the cells we allowed to regenerate for 5-10 minutes prior to mounting for TIRF microscopy.

To determine the swimming velocity, cells in fresh M medium were placed in a chambered plastic slide (Fisherbrand, 14-377-259) and observed using an inverted cell culture microscope (Nikon,TMS). Using a fixed exposure time of 1 s, images were taken using a MU500 camera (Amscope) and Topview software. The length of the trajectories resulting from the cells’ movements were analyzed using ImageJ and converted into μm/s.

### Probability of co-transport calculation

For our calculations, we assumed an IFT train frequency of 60/minute, which is close to that reported in serval studies of IFT in Chlamydomonas (Kozminski *et al*. 1993; Dentler 2005; Wingfield *et al*. 2017). Then, the probability of ARMC2-mS transport (P_(ARMC2-mS)_) in our data set is 0.268 (= 243 of 905 trains carried ARMC2-mS) and that for IDA3-NG transport (P_(IC2-NG)_) is 0.117. Multiplication of these two probabilities results in 0.03, as the expected probability (P_cotransport-calculated_) of ARMC2-mS and IDA3-NG being co-transport if they bind independently of each other to IFT trains. This value is close to 0.022, the observed probability of co-transports (P_cotransport-observed_). The probability of ARMC2-mS/RSP3-NG and ARMC2-mS/IC2-NG co-transports was calculated similarly.

## Abbreviations

CHX: cycloheximide
FP: fluorescent protein
IFT: intraflagellar transport
NG: mNeonGreen
mS: mScarlet-I
RS: radial spoke.

## Acknowledgements

We thank Yuqinq Hou and George Witman (UMASSMED) for sharing their proteomic data on ARMC2, Gui Zhang and Stephanie Chen for technical support, Win Sale (Emory University) for critical reading of the manuscript, and Paul Lefebvre and Matt Laudon (*Chlamydomonas* Resource Center) for the advice using on the CLiP library. This study was supported by a grant by the National Institutes of Health (R01GM110413 to K.L. and R015GM128130 to P.Y. and L.M.A.). The content is solely the responsibility of the authors and does not necessarily represent the official views of the National Institutes of Health. The authors declare that they have no conflict of interest.

## Competing interests

The authors declare that no competing interests exist.

## Supplementary materials

### Supplementary figures

**Figure 1 - Supplement 1.**
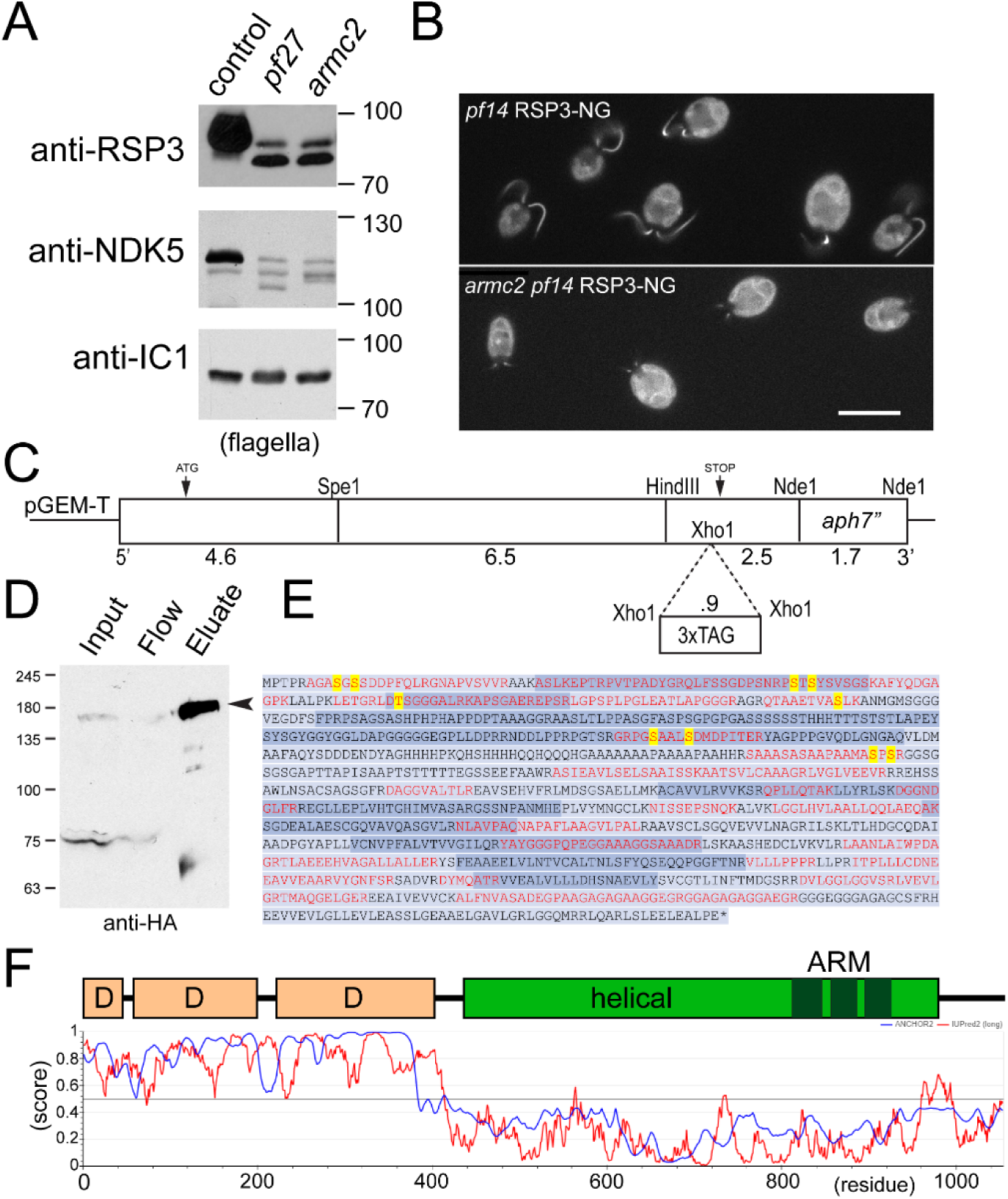
A) Western blots of flagella isolated from control, *pf27* and *armc2* probed with antibodies against RSP3 or RSP23/NDK5; to test for equal loading, we used antibodies directed against the outer dynein subunit IC1. In *pf27* and *armc2*, the RS proteins migrated faster due to altered phosphorylation and were less abundant. To visualize RSP3 and NDK5 isoforms, we used a 6% acrylamide gel and let it run until the 50 kD marker reached the gel front. B) In vivo epifluorescence imaging of *pf14* RSP3-NG rescue cells and *pf14 armc2* RSP3-NG cells. Bar = 10 μm. C) Description of the ARMC2 expression construct with or without a tag. The 15.3-kB genomic construct, was generated from ligating PCR-derived fragments of 4.6, 6.5, and 2.5 kB into the pGEM-T vector, along with the hygromycin resistance cassette (Hyg), using the indicated restriction sites. To insert the tags, an XhoI site was introduced into the 3’ fragment by fusing two PCR products. See Materials and Methods for details. D) Ni-NTA-chromatographic purification of ARMC2-3xTAG from whole cell extracts. The input, flow through (FLOW) and eluate were assessed by Western blot analysis using anti-HA. The ∼160 kD band enriched in the eluates (E) but absent in the flow-through was excised from a silver stained gel and subjected to mass spectrometry. E) Support of the 1053-residue ARMC2 sequence from mass spectrometry. Peptide sequences obtained by mass spectrometry of purified ARMC2-3xTAG and data mining are indicated in red letters. Yellow letters indicate the eight phosphorylation sites identified by phosphoproteomics of whole cell extracts (Wang *et al*. 2014). The encoding exons are marked in two shades of blue. F) Predicted structure of ARMC2. Top: D, disordered regions. In the helical region the three armadillo repeats (ARM) are indicated in dark green. Bottom: The N-terminal ∼400 residues of ARMC2 are predicted to be largely disordered (IUPred2, red line, score close to 1) and to assume a more structured configuration when binding to other proteins (blue line, ANCHOR2 score close to 1). The C-terminal region of ARMC2 is predicted to be largely ordered and encompasses three armadillo repeats (ARM).

**Figure 2 - figure supplement 1).**
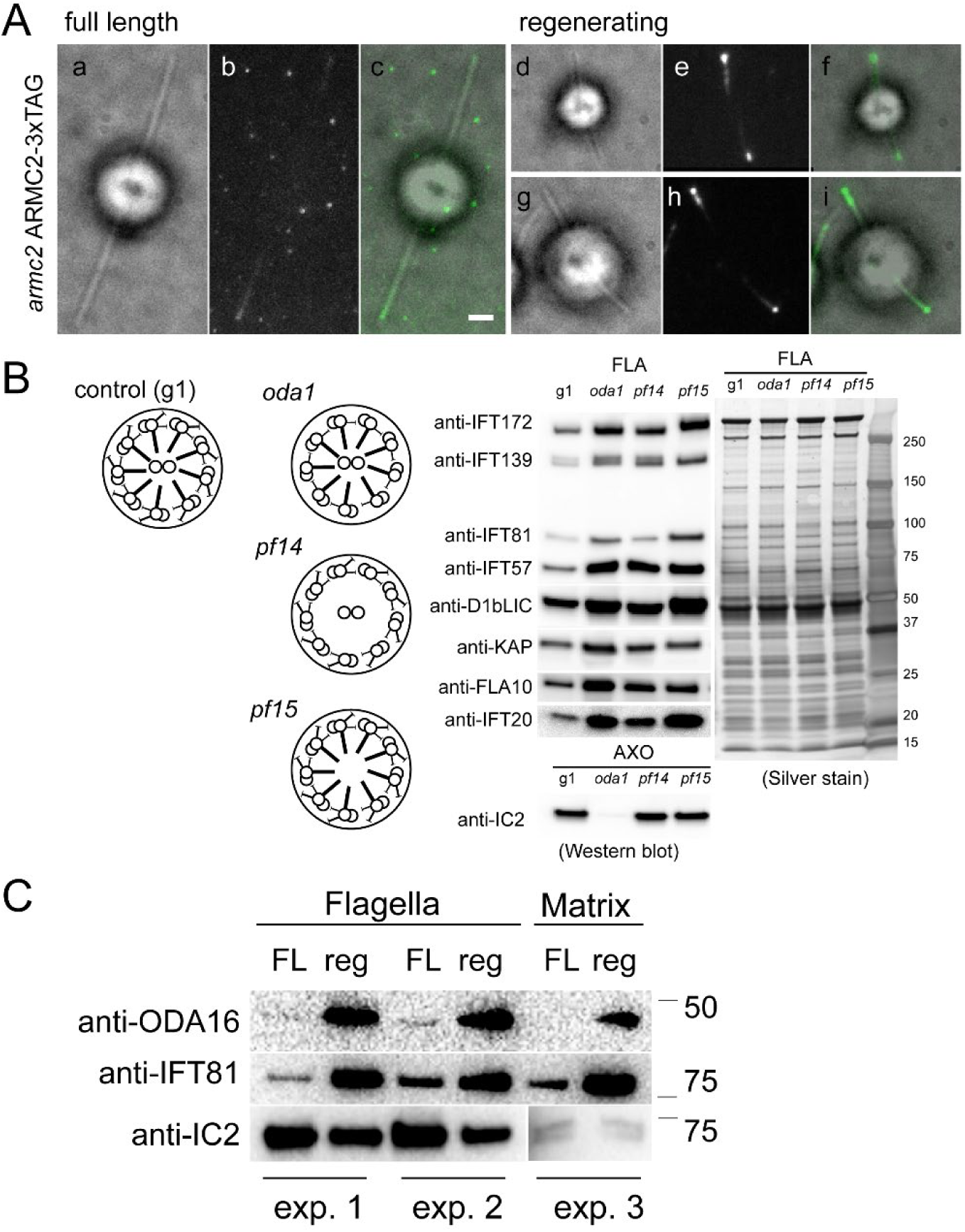
ARMC2-FP accumulates at the tip of growing flagella. A) Bright field (a, d, g), TIRF (b, e, h), and merged images (c, f, i) of ARMC2-3xTAG in full-length (a-c) and regenerating (d – i) flagella of the *armc2* ARMC2-3xTAG strain. Bar = 2 μm. B) Schematic presentation, western blot analysis and silver stained gel of isolated flagella from control (g1), and the *oda1, pf14* and *pf15* mutants. Note accumulation of IFT proteins in the flagella these motility mutants. C) Comparison of ODA16 levels in full-length (FL) and regenerating (reg) wild-type flagella from two independent biological replicates (exp. 1 and 2); from a third experiment (exp. 3) only the matrix fraction was available for analysis, explaining the weak signal for the axonemal protein IC2. For experiment 1 and 2, approximately equal number of flagella were loaded as apparent from the stronger IC2 bands in the FL samples.

**Figure 5 - figure supplement 1).**
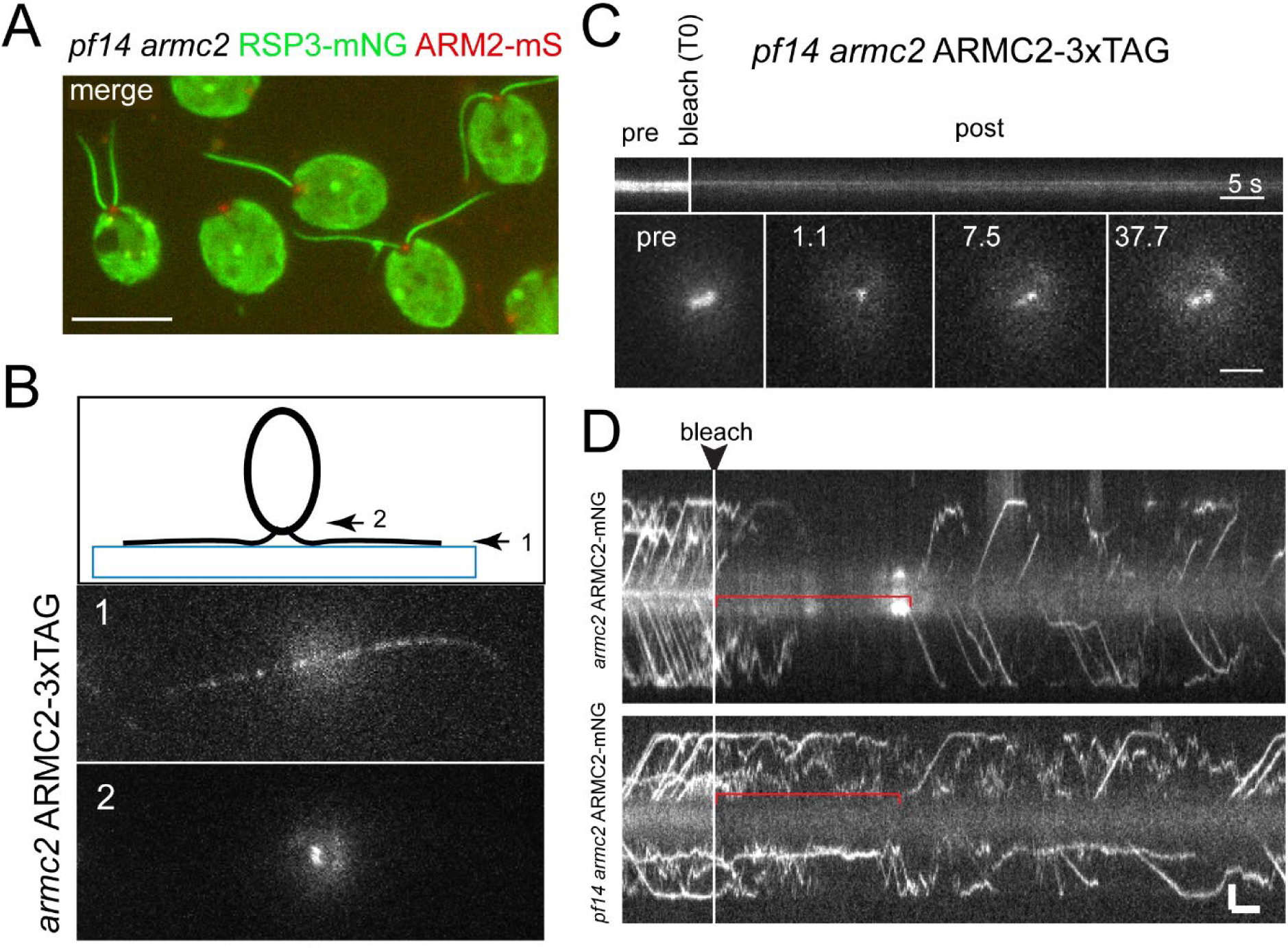
ARMC2-FP forms a pool near the basal bodies. A) 2-color epifluorescence image of the *pf14 armc2* RSP3-NG ARMC2-mS strain. Note accumulation of ARMC2-mS (red) at the flagellar base. Bar = 10 μm. B) Focal series using flat-angle “TIRF” illumination showing the flagella level (1) and the two ARMC2-3xTAG signals at the basal body level (2). The diagram illustrates the focal levels of the two images. C) FRAP experiment using a focused laser beam to bleach one of the two ARMC2-3xTAG signals at the flagella base. Note slow and partial recovery of the signal. Statistical analysis was not performed as only three cells were analyzed. Bars = 5 s and 2 μm. D) FRAP experiments to determine the dwell time of ARMC2-3xTAG in the basal body-associated pool. Regenerating flagella of the *armc2* ARMC2-3xTAG and the *pf14 armc2* ARMC2-3xTAG strains were analyzed. As in D, we used a focused laser beam to bleach the entire pool at the flagellar base and then analyzed the length of the gap before IFT of ARMC2-3xTAG resumed.

## Supplementary movies

**Supplementary Movie 1).**
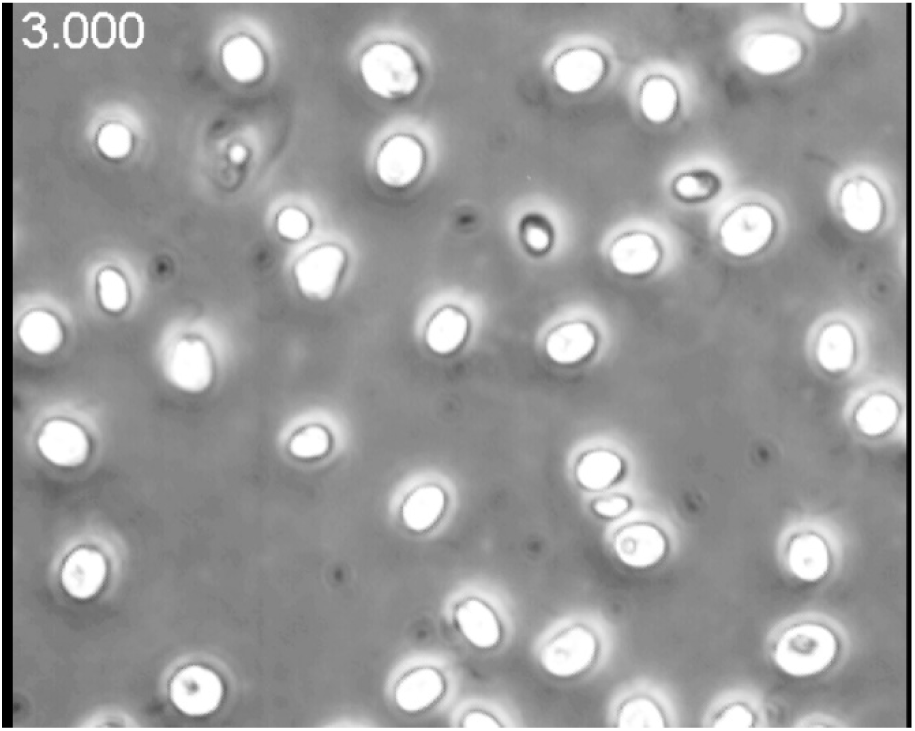
armc2 has a paralyzed flagella phenotype. Time lapse movie of armc2 mutant cells. The movie was recorded at 3 fps using phase contrast and a Nikon Eclipse 55i microscope equipped with a 40x/0.65 objective and a Zeiss AxioCam ERc5S. The timer displays seconds. The movie is related to Fig. 1B.

**Supplementary Movie 2).**
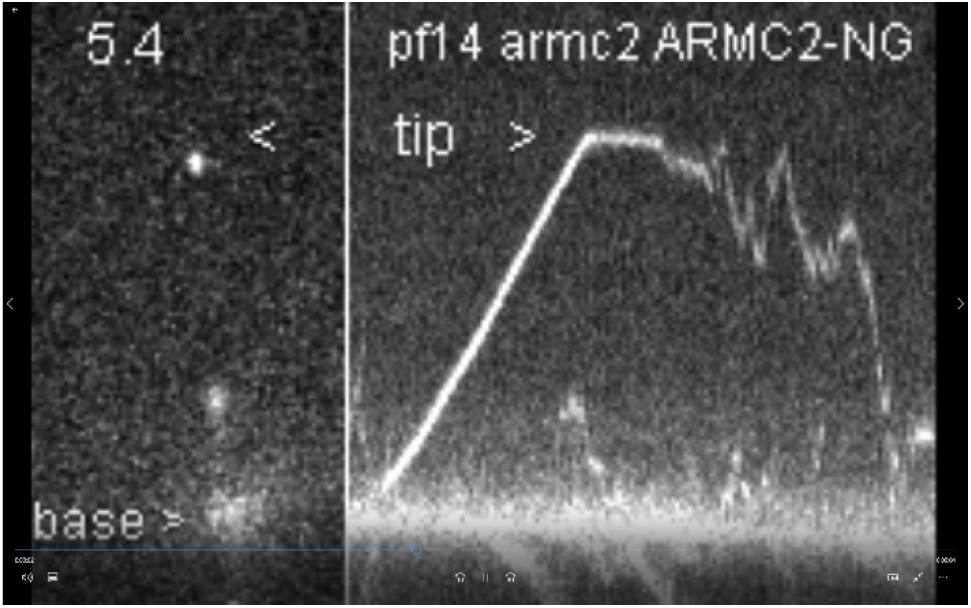
Anterograde IFT of ARMC2-3xTAG. In the ift54-2 2xDendra-IFT54 a subset of the fusion protein auto-converts from the green emitting into the red-emitting Dendra form allowing to visualizes individual IFT54 speckles during turnaround at the tip. The star marks the time points when Dendra^RED^-IFT54 speckles approach the tip. The corresponding kymograms are shown at the bottom. The movie was recorded at 20fps and the timer displays seconds. The movie is related to Fig. 2C.

**Supplementary Movie 3).**
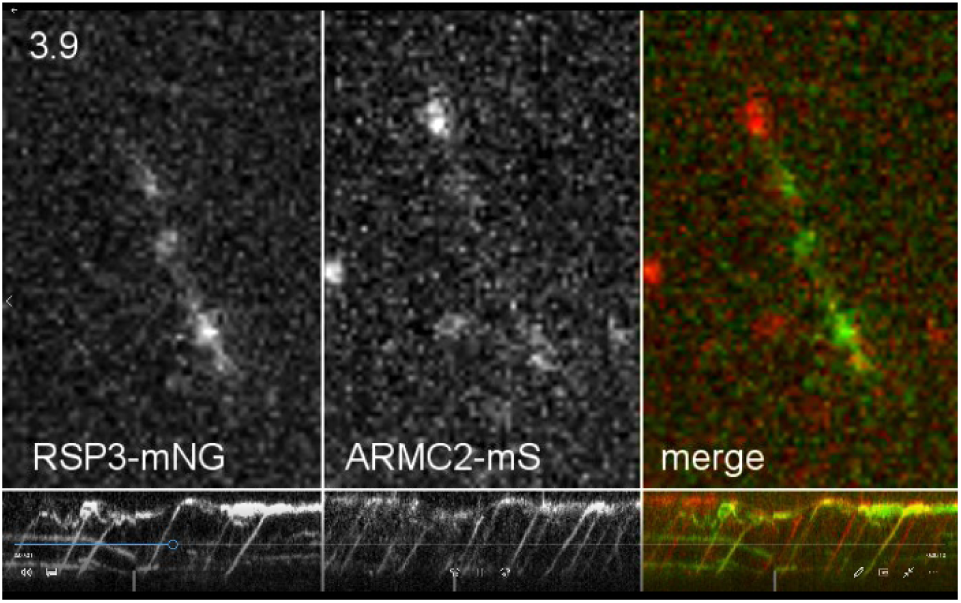
Co-transport of RSP3-NG and ARMC2-mS. Movies and corresponding kymograms of a flagellum form *armc2 pf14* ARMC2-mS RSP3-NG cells. Shown are the RSP3-NG and ARMC2 single channels and the merged image. An out-of-focus stretch of frames was deleted in the middle of the movie as indicated by white bars in the kymograms. The movie was recorded at 10 fps and the timer counts seconds. The movie is related to Fig. 3A.

**Supplementary Movie 4).**
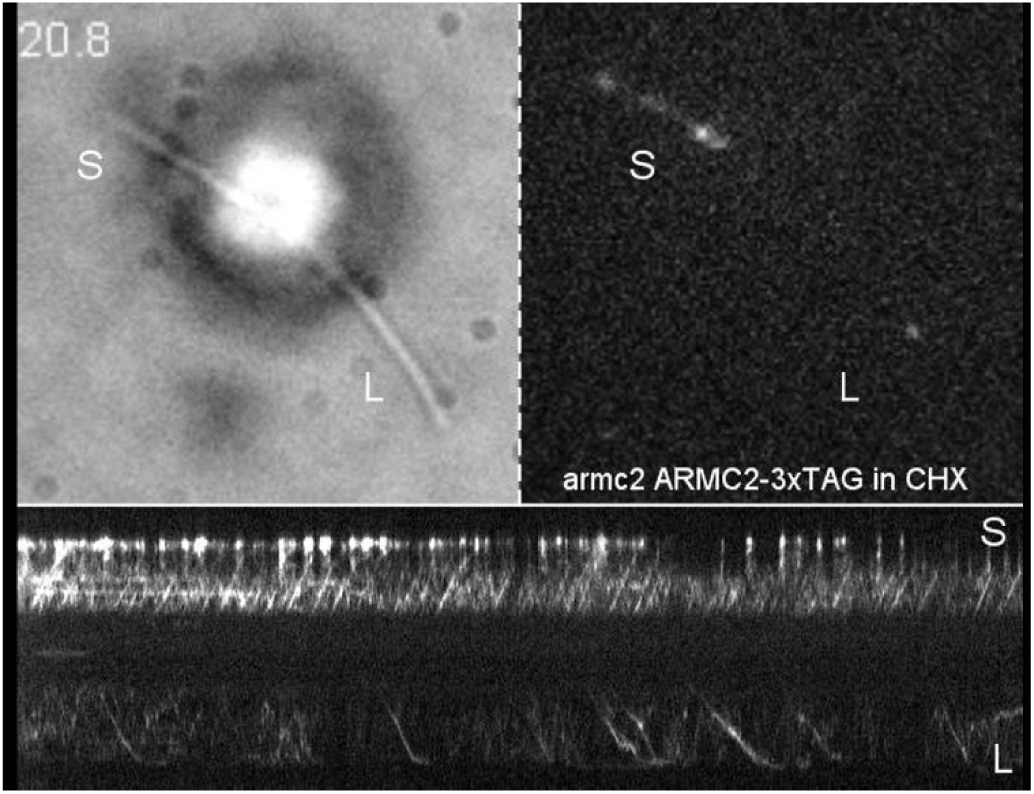
ARMC2-3xTAG transport in long-short cells. Bright field and TIRF video and the corresponding kymogram of a long-short *armc2* ARMC2-3xTAG cell. Cells were treated for one hour in CHX, sheared to generate long-zero cells and allowed to regrow missing flagella in the presence of CHX. The movie was recorded at 10 fps and the timer counts seconds. The movie is related to Fig. 5.

## Notes

### Competing Interest Statement

The authors have declared no competing interest.

